# CellTypeAI: Automated cell identification for scRNA-seq using local generative-AI

**DOI:** 10.64898/2026.03.03.709253

**Authors:** Rufus H. Daw, Harry R. Deijnen, Magnus Rattray, John R. Grainger

## Abstract

Single-cell RNA sequencing (scRNA-seq) cell annotation techniques rely on the matching of known defining marker genes to a given cell population. However, these methods may lack robustness to dynamic fluctuations in cell marker expression between patients, samples and pathologies. The advent of easy-to-implement predictive technologies, like generative-AI (gen-AI), has facilitated the introduction of computational workflows that improve otherwise inaccurate context-dependent cell type annotation. Here, we introduce CellTypeAI, a streamlined, scalable program developed for tissue context-dependent cell annotation of scRNA-seq datasets using modern gen-AI models, enhanced by retrieval augmented generation methods. Our implementation builds upon local gen-AI hosting technologies and directly integrates into scRNA-seq analysis pipelines. We show that CellTypeAI provides improved annotation accuracy compared to current conventional annotation methods and nascent cloud-based gen-AI approaches. As CellTypeAI leverages locally-run AI models, it can be applied to sensitive datasets, unlike approaches utilising online gen-AI tools such as ChatGPT, DeepSeek, or Claude. CellTypeAI presents a novel solution for tissue-specific cell type identification, overcoming traditional marker-based limitations via locally-deployed generative-AI models.

## Introduction

Accurate cell type annotation is a critical step in gathering meaningful insights from single-cell RNA sequencing (scRNA-seq) datasets. Manual ‘expert’ annotation of clusters is considered the current gold standard (Clarke et al., 2021). This is often a time-consuming and laborious task, given the ever-increasing sample sizes of scRNA-seq datasets, as sequencing costs continue to decline (Schwarze et al., 2020). Classically, computational solutions aiming to perform unbiased and accurate automated cell type annotation, in lieu of manual expert annotation, have relied on either gene marker-based (Ianevski et al., 2022), or reference-based methods (Domínguez Conde et al., 2022).

Both gene marker- and reference-based techniques typically use known ‘key marker genes’ of a given cell subset to explicitly, or implicitly, define and annotate cells within scRNA-seq datasets (Chen et al., 2024). To a degree, these methods are therefore reliant on consistent differential expression of ‘key marker genes’ for cell delineation and thus can be ignorant of the dynamic fluctuations in these genes between patients, samples, and pathologies (Rentzsch et al., 2025). Even when applied to simple biological contexts, the accuracy of current computational cell type annotation techniques can be low (Abdelaal et al., 2019). Moreover, the single-cell sequencing technologies that rely on computational cell identification have grown in scope, scale and complexity (Yu et al., 2022), underscoring the need to develop new computational methods that better support these increasingly complex and emerging technologies.

In recent years, generative artificial intelligence (gen-AI)-based predictive modelling techniques have improved in efficiency and accuracy (Awan et al., 2025). The advent of easy-to-implement predictive technologies, like gen-AI, enable their introduction into biological computational workflows. Such work includes efforts to develop generative pretrained transformers (GPTs) for cell type identification (Cui et al., 2024), and the integration of cloud-based large language models (LLMs) in single- and multi-prompt techniques for cell annotation (Hou & Ji, 2024; Ye et al., 2025). Importantly, these techniques have shown success in annotation accuracy against previous methods (Schaefer et al., 2025). However, the use of cloud-based LLMs for cell identification comes at a data security and financial cost (Burns et al., 2025). Similarly, reliance on, and integration of, these cloud-based LLMs into scRNA-seq annotation pipelines leaves users vulnerable to service availability issues, cost variability, reproducibility concerns, and version control issues (Jacobs & van Ginneken, 2019; Simkó et al., 2024). Thus, alternative methods of integrating LLMs into cell identification workflows need to be explored, for both data-sensitive applications, and the pursuit of reproducible, open-source methodologies for gen-AI-driven cell identification.

Since the transformer-based architecture was first introduced, the locally-deployed, open-source gen-AI field has developed alongside its cloud-based (and mostly closed-source) gen-AI counterparts, thus facilitating accessible, local deployment of small and large language models, with performance proportional to their parameter size and computational cost (Achiam et al., 2023; Touvron et al., 2023; Vaswani et al., 2017). Similarly, recent gen-AI integration techniques such as retrieval augmented generation (RAG) have significantly enhanced gen-AI output accuracy and efficacy through the injection of data additional to an LLM’s parameterised knowledge in any given prompt (Lewis et al., 2020).

Here, we introduce CellTypeAI, a streamlined, scalable program developed to perform context-dependent cell identification in scRNA-seq datasets, which leverages locally-run LLMs with enhanced RAG methods and optional ensemble methods. CellTypeAI builds upon local LLM hosting technologies and integrates directly into scanpy-based scRNA-seq analysis pipelines to enable accurate cell type identification of pre-clustered scRNA-seq data. We report that CellTypeAI shows improved cell identification accuracy compared to existing annotation methods.

## Methods

### 2.1 Dataset Collection

Five preprocessed scRNA-seq datasets were chosen for evaluation of CellTypeAI, using the previous manual expert annotation as the ground truth for cluster-by-cluster annotation evaluation.

#### 2.1.1 PBMC3K

The PBMC3K dataset was obtained directly from 10X Genomics (3K PBMCs v1.1.0 (https://support.10xgenomics.com/single-cell-gene-expression/datasets/1.1.0/pbmc3k). The expert-annotated cell clusters taken from scanpy’s “Preprocessing and clustering 3K PBMCs (legacy workflow)” were used as the ground truth for annotation accuracy calculation (Scanpy, 2025).

#### 2.1.2 Tabula Sapiens: Liver, Lung, and Small Intestine

The Tabular Sapiens datasets were obtained via the accession GSE201333. The expert-annotated cell clusters taken from the Tabular Sapiens: Liver, Lung, and Small Intestine h5ad.obs were used as ground truth for annotation accuracy calculation.

#### 2.1.3 Stroke PBMC

The Stroke PBMC dataset was generated in-house via peripheral blood mononuclear cells (PBMCs) taken from 2 stroke patients, at the in-patient stage, and at 6-9 months follow-up (**Supplementary Table 1, Supplementary Methods 1**). Approximately 33,000 cells per pool (8250 per individual, per stage) were loaded onto the 10x Chromium controller using Chromium Next GEM Single Cell 5’ Reagent Kits (10x Genomics) according to the manufacturer’s protocol. Sequencing was performed on an Illumina NovaSeq 6000 using a S1 flow cell, at ∼35,000 reads per cell. Data pre-processing was performed following scanpy’s standard vignette and clustering results can be found in **Supplementary Figure 1**. Manual cell annotation was provided by two independent immunologists at the Lydia Becker Institute of Immunology and Inflammation.

### 2.2 Model Selection

All models were hosted on Ollama (Ollama, Version 0.13.5). Three models were selected, based on their size, to represent different use scenarios. The first model was Phi4:14b, a 14B parameter small language model (SLM) developed by Microsoft and released in December 2024 (Abdin et al., 2024). Phi4:14b has a context window length of 16K. Because of its small size, Phi4:14b has a low RAM requirement (16GB), and was chosen to represent accuracy attainable when running on a personal laptop. The second model chosen was Qwen3:32b, developed by Alibaba, a 32B parameter dense model released August 2025 (Yang et al., 2025). Qwen3:32b has a 32K context length and medium RAM requirement (32GB). Qwen3:32b was selected to represent the accuracy attainable when running on a high-performance computer. The final model chosen was Qwen3:235b developed by Alibaba as part of the Qwen3 suite. This is a 235B parameter mixture of experts (MoE) LLM with a 32K context length. Because Qwen3:235b has a very high RAM requirement (250GB), this was used to represent accuracy attainable when running CellTypeAI on a high-performance compute cluster.

### 2.3 Evaluation of Models and Methods

All Ollama-hosted LLM runs were performed on a Mac Studio M3 Ultra using Metal Performance Shaders (32 core CPU, 80 core GPU, 512GB RAM). Google Gemini 2.5 Pro was used as the cloud-AI comparator (accessed September 2025) (Comanici et al., 2025). For both Ollama-hosted LLMs, and Gemini 2.5 Pro, a deterministic temperature was used (0.1), and reasoning/thinking modes were turned off where appropriate. The mean annotation accuracy was taken from three independent annotation ‘runs’ for local- and cloud-AI annotation runs. All were compared to the conventional annotation methods: ScType (version at commit hash 59a615e) and CellTypist (Version 1.7.1). The CellTypist models used for annotation were: Immune_All_Low (PBMC3K, Stroke PBMC), Healthy_Human_Liver (Tabular Sapiens: Liver), Human_Lung_Atlas (Tabular Sapiens: Lung), Cells_Intestinal_Tract (Tabular Sapiens: Small Intestine).

To evaluate all models and methods fairly, we developed a scoring system intended to penalise over- and under-specification in annotation but to allow for tissue-specific cell nomenclature. A range of 0-3 points were awarded per annotation based on the following criteria: 3 points for an exact or highly specific match (i.e. the model’s annotation was a precise match or a tissue specific equivalent), 2 points for the correct general category match (i.e. correct broader cell type but was either less or overly specific compared to the human annotation), 1 point for partially correct or related lineage match (i.e. the annotation was in a related lineage but not the correct final cell type); 0 points for a clear misclassification.

### 2.4 Data Availability

The Tabular Sapiens datasets can be obtained via the accession number: GSE201333. The PBMC3K dataset is publicly available from 10x Genomics (https://support.10xgenomics.com/single-cell-gene-expression/datasets/1.1.0/pbmc3k). Our Stroke PBMC dataset is available via the accession: E-MTAB-16654. The raw data obtained via CellTypeAI, and statistical analysis of this data, can be accessed via Zenodo (DOI: 10.5281/zenodo.18697111).

### 2.5 Code Availability

The source code for CellTypeAI is available on GitHub (https://github.com/rhdaw/celltypeai). This includes documentation for installation, and use. Additionally, the code used to process and analyse the Stroke PMBC dataset, CellTypeAI data acquisition, and manuscript graph generation is available in this GitHub repository.

## Results

### 3.1 Application architecture

CellTypeAI makes use of RAG, a prompt-engineering approach that uses a database to expand LLM prompts for improved cell type identification. CellTypeAI implements a user modifiable non-vectorised *cell_context* RAG database (JavaScript Object Notation (JSON) format, see **Figure 1a**). Within this *cell_context* database exists 16 different tissue reference datasets, with up to 35 different cell types and their defining genes per tissue. As part of the RAG system, these genes are retrieved when making a call to CellTypeAI as additional reference material to aid prompt response accuracy and thus cell annotation. As *cell_context* is non-vectorised, users can add and remove tissues, cell types, and defining genes to their specification in the pursuit of improved cell annotation.

**Figure 1.**
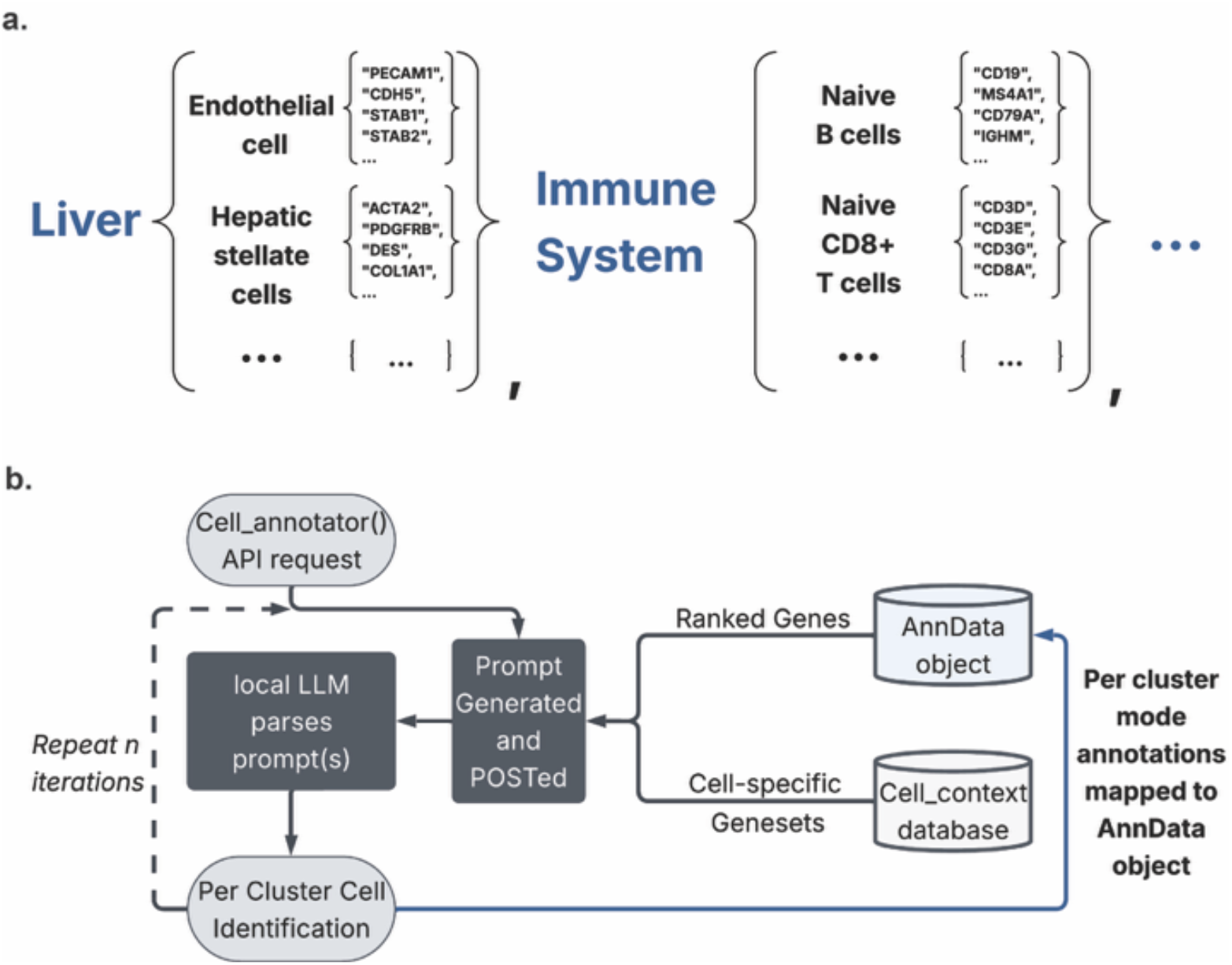
Visual representation of the CellTypeAI API and RAG database. **a**, A graphical representation of the *cell_context* JSON architecture whereby cell types and their corresponding marker genes are grouped by tissue type. ‘…’ denote continuations of *cell_context*’s nested JSON structure. **b**, A sequence diagram of the CellTypeAI Application Programming Interface (API). Arrows denote the step-by-step API flow, dotted arrows denote optional flow based on the *n_iterations* argument, the blue arrow denotes the final logic step of per cluster annotation.

Our initial *cell_context* database, and the ‘tissue: cell type: gene-set’ combinations therein, was constructed via comprehensive assessment of the current literature on a tissue-by-tissue, cell type by cell type basis. Although the *cell_context* database is not exhaustive, it includes most cell types within each tissue. Moreover, our intention here was not to create a comprehensive database but rather provide a functional starting point and framework for further iteration and addition by domain experts. A full table detailing all tissue: cell type: gene-set combinations can be found in **Supplementary Table 2**.

CellTypeAI builds upon Ollama’s REST Application Programming Interface (API) to host local open-source LLMs, and parse scRNA-seq information to them, to obtain cell cluster annotations (**Figure 1b**). The user initialises CellTypeAI via the *cell_annotator* function, taking the AnnData object, ‘tissue’ type, number of iterations (ensemble methods), the number of genes to include in the prompt, and the species as arguments. Per cluster, ranked genes (calculated by scanpy’s *scanpy.tl.rank_genes_groups*) are inserted into the engineered prompt. Based on *cell_annotator*’s tissue argument, a series of tissue-specific cell types and their defining genes are inserted into the prompt as reference material, as part of the RAG system. From here, the finalised prompt is parsed by the Ollama-hosted LLM and an annotation is returned. This process is repeated per cluster. If ensemble methods are chosen, this process is then repeated for *n_iterations* and the mode annotation is mapped back to its respective cluster. If ensemble methods are not chosen, this process is skipped and the first (and only) LLM-based annotations are mapped instead.

Ollama has an extensive and continuously expanding library of publicly available LLMs. Here, we evaluated CellTypeAI using three separate open-source models, intentionally picked to represent annotation accuracy attainable in specific computationally restrained scenarios. Using these models, we systematically assessed CellTypeAI’s ability to annotate five human scRNA-seq datasets. The CellTypist (Domínguez Conde et al., 2022) and ScType (Ianevski et al., 2022) methods were also implemented for comparison. Because CellTypeAI generates an engineered prompt (itself parsed to the locally-run LLM), we extracted this prompt and parsed it to the cloud-based LLM Google Gemini 2.5 Pro for evaluation and comparison of cloud-based LLMs (Comanici et al., 2025). All cell type annotations via models and methods were evaluated against a ground truth set by the manual cluster annotations accompanying each dataset. The annotation accuracy of all methods and models was measured using an accuracy scoring system (See Methods: **Evaluation of Models and Methods**).

### 3.2 Overall CellTypeAI performance

Interrogation of the PBMC3K dataset annotations showed that CellTypeAI has excellent max mean annotation accuracy when running Qwen3:235b (94.4% ± 6.4) compared to cloud-AI-based solutions (Gemini 2.5 Pro: 86.6% ± 2.4), and conventional cell annotation methods (CellTypist: 70.8% ± 0, ScType: 54.2% ± 0) (**Figures 2a, 2b**). Notably, increased model size led to an increase in max mean annotation accuracy, with Qwen3:235b showing the best performance, followed by Qwen3:32b (87.5% ± 0), and Phi4:14b (70.8% ± 0). Qwen3:32b similarly outperformed Gemini 2.5 Pro, CellTypist, and ScType, while Phi4:14b showed consistent but limited performance, outperforming ScType, but only just reaching the same max mean annotation accuracy of CellTypist. Ensemble methods appeared to provide little benefit to annotation accuracy, with no obvious relationship between prompt iterations per run and max mean annotation accuracy.

**Figure 2.**
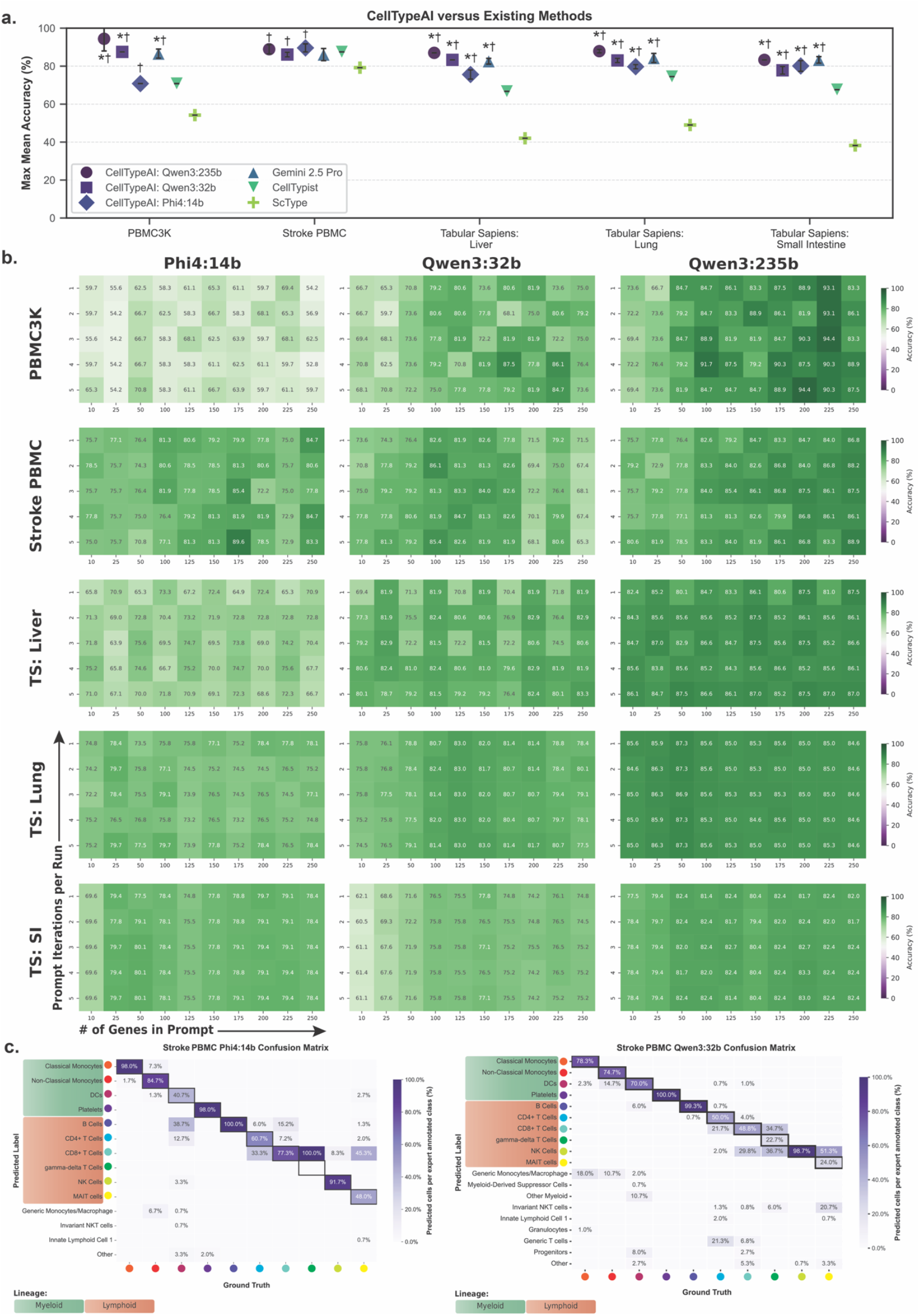
Assessment of CellTypeAI driven annotation illustrates its improved annotation accuracy compared to conventional identification methods. **a**, Comparison of the maximum mean annotation accuracy achieved across all annotation runs (comprised of a ‘prompt iteration / genes in prompt’ combination), per scRNA-seq dataset. * and † denote a significant difference in annotation accuracy of a given CellTypeAI:model combination versus CellTypist or ScType, respectively (see **Supplementary Table 3** for data on degrees of significance). Error bars denote standard deviation. n = 3. **b**, Heatmaps denoting average annotation accuracies across all CellTypeAI:model combinations, and datasets, across a range of ‘prompt iteration / genes in prompt’ combinations (TS = Tabular Sapiens, SI = Small Intestine), n = 3. **c**, Confusion matrices denoting the column normalized percentage overlap of expert-annotated cell types (i.e. Ground Truth) versus the CellTypeAI: Phi4:14b (left) and CellTypeAI: Qwen3:32b (right) annotations. Unmarked predicted labels refer to predicted labels generated by CellTypeAI that do not match the Ground Truth category.

The Tabular Sapiens: Liver, Lung and Small Intestine datasets were chosen to determine CellTypeAI’s accuracy in complex tissue scenarios. Across these datasets, there was a positive relationship between model size and max mean annotation accuracy, with little to no improvement when implementing ensemble methods (**Figures 2a, 2b**). CellTypeAI running Qwen3:235b had the highest max mean annotation accuracy across all three tissue datasets (Liver: 87.0% ± 0, Lung: 87.9% ± 0.9, Small Intestine: 83.3% ± 0) when compared to all models and methods. Notably, almost all CellTypeAI:model combinations had significantly improved annotation accuracy compared to ScType and CellTypist (**Supplementary Table 3**) in the Tabular Sapiens and PBMC3K datasets. Like in PBMC3K, CellTypeAI running Qwen3:235b or Qwen3:32b exhibited comparable annotation accuracy compared to Gemini 2.5 Pro, and outperformed CellTypeAI running Phi4:14b throughout the Tabular Sapiens datasets.

An in-house sequenced stroke PBMC dataset was included to assess CellTypeAI’s accuracy in a disease dataset, and to explore the effect of potential data leakage on annotation accuracy. Both PBMC3K and the Tabular Sapiens datasets were made publicly available prior to Phi4 and Qwen3’s training cut-off dates (Abdin et al., 2024; Yang et al., 2025), thus both datasets run the risk of having been incorporated into Phi4 and Qwen3’s training and development. Conversely, our Stroke PBMC scRNA-seq dataset has remained internal until now and cannot have been included in the training data used to build any of the LLMs used here. In this dataset, ScType showed the lowest annotation accuracy (79.2% ± 0), whilst Phi4:14b had the highest (89.6% ± 2.1) (**Figures 2a, 2b**). Surprisingly, Phi4:14b outperformed the significantly larger Qwen3:235b (88.9% ± 2.4), Qwen3:32b (86.1% ± 1.2), and the cloud-based Gemini 2.5 Pro (84.0% ± 3.2). CellTypeAI-based annotation was either marginally more accurate than, or equivalent to, current cloud-based AI and conventional annotation methods on this dataset.

### 3.3 Annotation prediction across models

To better understand why Qwen3:32b showed worse annotation accuracy in the Stroke PBMC dataset, as compared to Phi4:14b, we generated confusion matrices for both models (**Figure 2c**). Here, Qwen3:32b showed poorer cell type prediction across monocyte lineages (Classical Monocytes: 78.3%, Non-Classical Monocytes: 74.7%) and most T cell lineages (CD4^+^ T cells: 50%, CD8^+^ T cells 48.8%, MAIT cells: 24%), compared to Phi4:14b (Classical Monocytes: 98.0%, Non-Classical Monocytes: 84.7%, CD4^+^ T cells: 60.7%, CD8^+^ T cells 77.3%, MAIT cells: 48%). However, Phi4:14b failed to correctly annotate gamma-delta T cells, instead opting to classify them as CD8^+^ T cells. We hypothesise that differences in parameterised knowledge and training techniques between the Phi4 and Qwen3 models may have contributed to Qwen3:32b’s poorer prediction of monocyte and T cell subsets in this dataset, potentially overcome by Qwen3:235b’s improved architecture and model size (Yang et al., 2025). Thus, model selection, alongside model size, may contribute to CellTypeAI’s annotation performance.

From our previous data, it was apparent that CellTypeAI running Qwen3:235b offered the greatest annotation accuracy of the CellTypeAI:model combinations. As an evaluation of the best-case scenario, we further assessed Qwen3:235b’s annotation performance across the five datasets via a cumulative confusion matrix (**Figure 3a**). Across all datasets, CellTypeAI: Qwen3:235b excelled at annotating most lymphoid, myeloid and non-immune cell subsets. However, over-specific annotation was observed in instances where the ground truth annotations were generic. For example, the majority of ‘T Cell’ or ‘Myeloid Cell’ ground truth annotations were over-specified to particular T cell subsets or monocyte / macrophage subsets, respectively. Despite this, these over-specific annotations remained within the general lineage categories (i.e. myeloid, lymphoid) of the ground truth label. Similarly, instances of misclassification (gamma-delta T cells, CD4^+^ and CD8^+^ T Cells) or under-specification (Hepatic Stellate Cells, Fibroblasts) by Qwen3:235b mostly occurred between closely related lineages, and within general lineage categories (lymphoid and non-immune, respectively).

**Figure 3.**
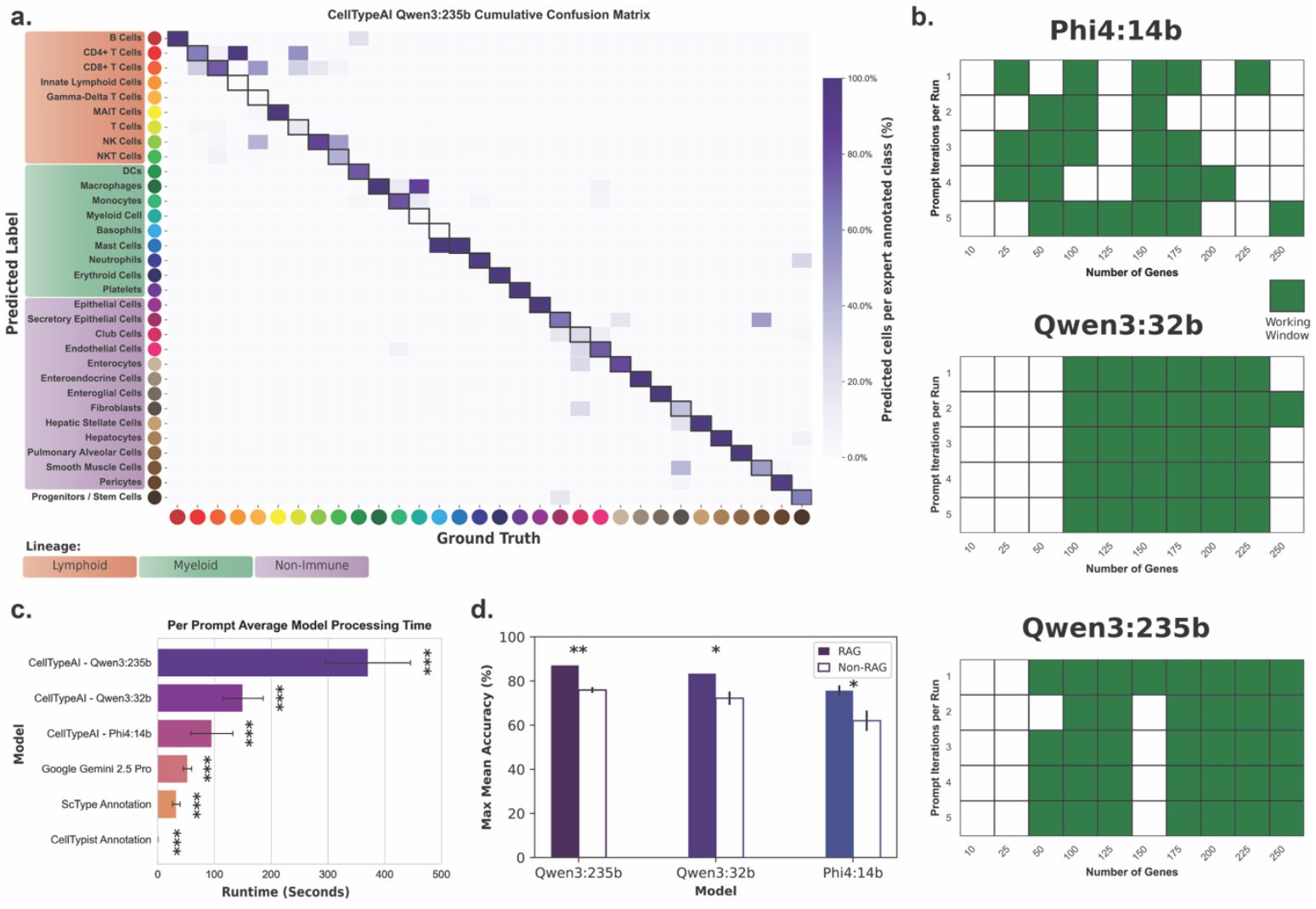
Benchmarking of CellTypeAI across model combinations. **a**, A cumulative confusion matrix compiled from the major annotation labels (split by lineage), generated by CellTypeAI: Qwen3:235b across the PBMC3K, Stroke PBMC, and Tabular Sapiens datasets used in this paper. Purple shading denotes column normalised percentage overlap of expert-annotated cell types. **b**, ‘Working window’ of prompt iteration / genes in prompt combinations most likely to give the user an annotation with accuracy above the average attainable with that CellTypeAI:model combination, denoted by green shading. **c**, The mean model processing wall-clock time per single prompt, computed across all single annotation iteration runs on the PBMC3K dataset, indicative of the proportional time complexity of different CellTypeAI:model combinations and current annotation methods. *** = p < 0.001. n = 30 per CellTypeAI:model combination. n =10 for Google Gemini 2.5 Pro (time shown is ‘API latency’), ScType, CellTypist. Error bars denote standard deviation. **d**, The impact of RAG techniques on max mean annotation accuracy on the Tabular Sapiens: Liver dataset, across all three CellTypeAI:model combinations. Shaded bars denote a RAG approach; clear bars denote a RAG-less approach. Error bars denote standard deviation. * p < 0.05 | ** = p < 0.01, n = 3.

### 3.4 Computational benchmarking

With several prompt iteration / gene combinations available to the user, we next analysed our annotation results across all five datasets to determine the most effective iteration / gene combinations per model. Min-max normalisation of mean annotation accuracy of each iteration / gene combination was used to indicate a CellTypeAI ‘working window’ in which the highest annotation accuracy can be found (**Figure 3b**). Here, normalised values greater than the normalised mean of that model across all five datasets were included in the working window. Best accuracy in both Qwen3 models could be found in iteration / gene combinations where the number of genes in the prompt was greater than 100. Across both Qwen3 models, prompt iterations per run had minimal effect on whether an iteration / gene combination was included in the working window. Phi4:14b had a sparser working window, with peak accuracy attainable at over 150 genes in prompt. Qwen3 showed the most stable working window, likely attributed to its model size and context window length (32K tokens), markedly greater than Phi4:14b (16K) (Abdin et al., 2024; Yang et al., 2025).

Given the computational overhead of LLM inference, we wanted to benchmark CellTypeAI’s run time in comparison to other methods. There was a positive association between model size and processing time across all CellTypeAI hosted models (**Figure 3c**). Across all model combinations, CellTypeAI had longer processing times compared to Gemini 2.5 Pro, ScType and CellTypist. Despite this, on our system CellTypeAI per prompt model processing time took, at most, 8 minutes to complete when annotating the PBMC3K dataset. However, as our testing was limited to Metal Performance Shaders, rather than the more efficient Compute Unified Device Architecture (CUDA), we would suggest that users using CellTypeAI in CUDA supported scenarios will see significant improvements in processing time (Oh & Jung, 2004).

Lastly, as RAG is a central component of CellTypeAI, we wished to assess CellTypeAI in the absence of RAG, to understand its impact on annotation accuracy (**Figure 3d**). Here, CellTypeAI was run as described in **Figure 1a**, but the *cell_context* JSON no longer supplied supplementary cell marker information to the engineered prompt. Limiting this experiment to only the Tabular Sapiens: Liver dataset, we observed a 10–15% increase in annotation accuracy across all three models, indicating a small but significant uplift in annotation accuracy when RAG techniques are employed.

## Discussion

Accurate scRNA-seq cell annotation remains a complex issue, with current methods prone to misclassification in tissue scenarios and disease states (Abdelaal et al., 2019). LLM-based annotation methods, such as scExtract and CellWhisperer, show promise over conventional methods, given their ability to integrate additional information into their classification pipelines (Schaefer et al., 2025; Wu & Tang, 2025). Here, we show that CellTypeAI, a local LLM-driven cell annotation program, provides improved annotation accuracy compared to current conventional methods and nascent cloud-based LLM approaches. As CellTypeAI is run locally, be it on the user’s own computer or an accessible high performance compute cluster, this comes at no additional cost, unlike cloud-AI approaches which incur API access charges (Burns et al., 2025).

CellTypeAI is augmented by a user modifiable RAG database, stored in JSON format. Whilst the current *cell_context* database is extensive enough to provide good annotation results, users are also able to add or remove cell types, new tissues, and disease state contexts, to improve annotation accuracy in their use cases. Most importantly, of the few LLM-based cell annotation programs published, most rely on cloud-based LLMs for their annotation (Cui et al., 2024; Wu & Tang, 2025). Although this removes the need for local compute, it comes with serious data security concerns. CellTypeAI avoids these issues by using local models, enabling reproducibility, optimisation, and secure use with sensitive datasets.

The local-LLM research community continues to develop an expanding library of computationally capable LLMs. As CellTypeAI leverages Ollama’s REST API, acting as a framework for local-LLM driven cell identification, any current or future open-source model made available on Ollama can be used for cell annotation purposes via CellTypeAI. This design choice also enables CellTypeAI to run across systems with varying compute resources, with annotation accuracy scaling accordingly. We noted that even when running the smallest model (Phi4:14b), annotation accuracy was often on par with, or slightly better than, current conventional methods. Despite this, compute availability remains a major obstacle for accurate cell annotation. One solution to this is the implementation of quantised LLMs, which reduce computational requirements of high-parameter models with marginal impact on inference accuracy (Li et al., 2024). Their applicability was not assessed in this study, but we suggest that weight or key-value cache quantised models may offer improved annotation accuracy on systems with weaker computational capabilities.

## Author Contributions

Rufus H. Daw (Conceptualisation, Data curation, Investigation, Funding acquisition, Methodology, Software, Visualisation, Writing—original draft, Writing—review & editing), Harry R. Deijnen (Investigation, Writing— original draft, Writing—review & editing), Magnus Rattray (Conceptualisation, Funding acquisition, Supervision, Writing— review & editing), John R. Grainger (Funding acquisition, Supervision, Writing—review & editing).

