## Supplementary Information for "CellTypeAI: Automated cell identification for scRNA-seq using local generative-AI"

| Supplementary Table 1. Patient data table for Stroke PBMC dataset. | | |
|---|---|---|
| **Study_ID** | 4133-066 | 4133-079 |
| **gender_usuk_label** | Male | Male |
| **Age_years** | 60 | 72 |
| **Ethnicity** | White | White |
| **NIHSS** | 7 | 9 |
| **Date_of_Enrollment** | 44665 | 44706 |
| **Time_of_symptom_onset** | 44663.41667 | 44705.91667 |
| **Time_symptoms_were_recognized** | NA | NA |
| **Time_last_known_to_be_neurologically_normal_(LSN/LKW)** | NA | NA |
| **Date/Time_Blood_Drawn_** | 44665.40972 | 44706.42708 |
| **blooddrawdiff** | 47.833333 | 12.25 |
| **Lesion_Location** | Left | Left |
| **Lacunar** | No | Unknown |
| **IV_Thrombolysis** | No | Yes |
| **Was_intra-arterial_thrombectomy_undertaken__?** | No | No |
| **stroke_AMPM** | AM | PM |
| **stroke_time_of_day** | 2 | 4 |
| **Is_ECG_available?_** | Yes | Yes |
| **Left_Ventricular_Hypertrophy?** | No | No |
| **Atrial_Fibrillation_(Afib)?** | No | No |

**Supplementary Methods 1: Stroke PBMC Processing**

As part of the Stroke-Immune Pathways and Cognitive Trajectory (Stroke-IMPaCT) Study, venous blood was drawn from ischaemic stroke patients within 96 h of stroke symptom onset or time last seen well, as well as at the 6-9-month follow-up timepoint. Blood was collected in ethylenediaminetetraacetic acid (EDTA) tubes and processed within 4 h.

To obtain peripheral blood mononuclear cells (PBMCs), EDTA blood was diluted 1:1 with phosphate-buffered saline (PBS) supplemented with 2% foetal bovine serum (FBS) and centrifuged at 1200 g for 15 min at 20°C in SepMate™ PBMC Isolation Tubes (StemCell Technologies, Vancouver, Canada) with Ficoll-Paque™ PLUS density gradient media (Cat. No. 17144003; Cytiva Life Sciences, Amersham, UK). Following centrifugation, the top supernatant layer containing PBMCs was transferred to 50 mL falcon tubes, diluted with 20 mL PBS + 2% FBS and centrifuged at 500 g for 10 min at 20°C. Supernatant was discarded, and the cell pellet was resuspended in ACK Lysing Buffer (Cat. No. A1049201; Gibco, Waltham, USA) for 5 min at room temperature. PBS (10 mL) + 2% FBS was added and tubes centrifuged at 500 g for 5 min at 20°C. The cell pellet was resuspended in 20 mL FBS + 2% FBS and cells were counted before centrifugation at 500 g for 10 min at 20°C. Cells were resuspended at a final density of 5-6 × 106 cells / 500 µL CryoStor® CS10 Freezing Medium (Cat. No. 100-1061; StemCell Technologies), and frozen at -80°C in CoolCell™ Cell Freezing Vial Containers (Cat. No. 15542771; Corning, New York, USA). Finally, PBMC aliquots were transferred to liquid nitrogen until ready for use.

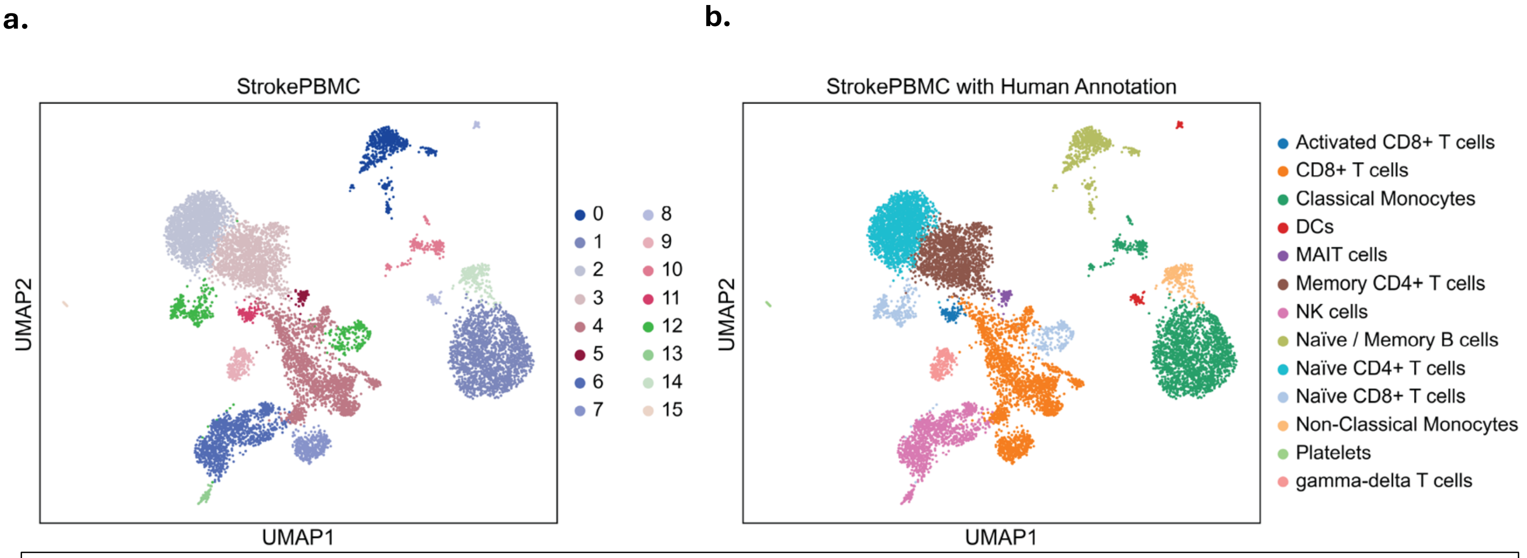

**Supplementary Figure 1. UMAP visualization of Stroke PBMC dataset. a**, UMAP visualization of leiden (0.3 resolution) clustering of Stroke PBMC single cell RNA sequencing samples. **b**, UMAP visualization of Stroke PBMC dataset with consensus human annotation of clusters depicted in **a**.

**Supplementary Table 2. Contents of cell_context.json in tabular form, listing all tissues, cell types, and their defining marker genes.**

| Tissue | Cell Type | Markers | Tissue | Cell Type | Markers |
|---|---|---|---|---|---|
| Immune system | Pro-B cells | CD19,CD79A,MS4A1,VPREB1,IGLL1,DNTT,RAG1,RAG2,CD24,KIT,IL7R | Intestine | Eosinophils | CCR3,IL5RA,SIGLEC8,EPX,RNASE2,RNASE3,PRG2,CLC,HPGDS,PTGDR2 |
| Immune system | Pre-B cells | CD19,CD79A,MS4A1,VPREB3,IGLL1,CD24,IL7R,PAX5,EBF1 | Intestine | Basophils | FCER1A,CD200R3,ENPP3,IL3RA,CD33,TPSAB1,TPSB2,HDC,GATA2 |
| Immune system | Naive B cells | CD19,MS4A1,CD79A,IGHM,IGHD,CD27,SELL,IL4R,CCR7,BACH2 | Intestine | Mast cells | KIT,FCER1A,CPA3,TPSAB1,TPSB2,MS4A2,ENPP3,CD63,CD9 |
| Immune system | Memory B cells | CD19,MS4A1,CD79A,CD27,CD38,SDC1,IGHG1,IGHG2,IGHG3,IGHG4,IGHA1,IGHA2,AICDA,BANK1,PAX5 | Intestine | Naive B cells | CD19,MS4A1,CD79A,IGHM,IGHD,CD27,SELL,IL4R,CCR7,BACH2 |
| Immune system | Plasma B cells | SDC1,CD38,XBP1,PRDM1,SLAMF7,DERL3,MZB1,IGHG1,IGHG2,IGHG3,IGHG4,IGHA1,IGHA2,IGLC2,IGKC | Intestine | Memory B cells | CD19,MS4A1,CD79A,CD27,CD38,SDC1,IGHG1,IGHG2,IGHG3,IGHG4,IGHA1,IGHA2,AICDA,BANK1,PAX5 |
| Immune system | Naive CD8+ T cells | CD3D,CD3E,CD3G,CD8A,CD8B,CCR7,SELL,LEF1,TCF7,IL7R,CD27,CD45RA | Intestine | Plasma B cells | SDC1,CD38,XBP1,PRDM1,SLAMF7,DERL3,MZB1,IGHG1,IGHG2,IGHG3,IGHG4,IGHA1,IGHA2,IGLC2,IGKC |
| Immune system | Naive CD4+ T cells | CD3D,CD3E,CD3G,CD4,CCR7,SELL,LEF1,TCF7,IL7R,CD27,CD45RA,FOXP1 | Intestine | Naive CD8+ T cells | CD3D,CD3E,CD3G,CD8A,CD8B,CCR7,SELL,LEF1,TCF7,IL7R,CD27,CD45RA |
| Immune system | Memory CD8+ T cells | CD3D,CD3E,CD3G,CD8A,CD8B,CD44,IL7R,CD69,EOMES,TBX21,IFNG,GZMK,GZMB,CD45RO | Intestine | Naive CD4+ T cells | CD3D,CD3E,CD3G,CD4,CCR7,SELL,LEF1,TCF7,IL7R,CD27,CD45RA,FOXP1 |
| Immune system | Memory CD4+ T cells | CD3D,CD3E,CD3G,CD4,CD44,IL7R,CD69,ICOS,CCR5,CCR6,CXCR3,TBX21,GATA3,RORC,FOXP3,CD45RO | Intestine | Memory CD8+ T cells | CD3D,CD3E,CD3G,CD8A,CD8B,CD44,IL7R,CD69,EOMES,TBX21,IFNG,GZMK,GZMB,CD45RO |
| Immune system | Effector CD8+ T cells | CD3D,CD3E,CD3G,CD8A,CD8B,GZMB,PRF1,IFNG,KLRG1,CX3CR1,TBX21,EOMES,CCL5,CD27,SELL | Intestine | Memory CD4+ T cells | CD3D,CD3E,CD3G,CD4,CD44,IL7R,CD69,ICOS,CCR5,CCR6,CXCR3,TBX21,CD45RO |
| Immune system | Effector CD4+ T cells | CD3D,CD3E,CD3G,CD4,IFNG,TNF,IL2,IL4,IL17A,IL21,ICOS,CD40LG,TBX21,GATA3,RORC,BCL6,PRDM1 | Intestine | Effector CD8+ T cells | CD3D,CD3E,CD3G,CD8A,CD8B,GZMB,PRF1,IFNG,KLRG1,CX3CR1,TBX21,EOMES,CCL5,CD27,SELL |
| Immune system | CD4+ T regulatory cells | FOXP3,IL2RA,CTLA4,IKZF2,TNFRSF18,CD4,PTPRC,LAG3 | Intestine | Effector CD4+ T cells | CD3D,CD3E,CD3G,CD4,IFNG,TNF,IL2,IL4,IL17A,IL21,ICOS,CD40LG,TBX21,GATA3,RORC,BCL6,PRDM1 |
| Immune system | γδ-T cells | CD3D,CD3E,CD3G,TRDC,TRGC1,TRGC2,NKG7,KLRB1,KLRG1,CCL5,CD160 | Intestine | γδ-T cells | CD3D,CD3E,CD3G,TRDC,TRGC1,TRGC2,NKG7,KLRB1,KLRG1,CCL5,CD160 |
| Immune system | MAIT cells | PTPRC,CD3D,SLC4A10,KLRB1,TRAV1-2,IL18R1,CCR6,RORC,ZBTB16 | Intestine | CD8+ NKT-like cells | CD3D,CD8A,CD8B,NCAM1,NKG7,KLRB1,KLRD1,GZMB,PRF1,TBX21,EOMES,SLAMF7 |
| Immune system | Follicular Helper T cells | PTPRC,CD4,BCL6,CXCR5,PDCD1,ICOS,SLAMF6,CD40LG,MAF,ASCL2 | Intestine | CD4+ NKT-like cells | CD3D,CD4,NCAM1,NKG7,KLRB1,GZMK,TBX21,EOMES,SLAMF6 |
| Immune system | Invariant NKT cells | PTPRC,CD3D,ZBTB16,TRAV10,TRBV25,SLAMF1,SLAMF6,CD4,CD8A | Intestine | Natural killer cells | NCAM1,NKG7,KLRF1,KLRD1,KLRB1,KLRC1,KLRC2,FCGR3A,GZMB,PRF1,EOMES,TBX21 |
| Immune system | CD8+ NKT-like cells | CD3D,CD8A,CD8B,NCAM1,NKG7,KLRB1,KLRD1,GZMB,PRF1,TBX21,EOMES,SLAMF7 | Intestine | Platelets | ITGA2B,ITGB3,GP1BA,GP9,PPBP,PF4,SELP,TUBB1,CD9 |
| Immune system | CD4+ NKT-like cells | CD3D,CD4,NCAM1,NKG7,KLRB1,GZMK,TBX21,EOMES,SLAMF6 | Intestine | Megakaryocyte | ITGA2B,ITGB3,GP1BA,GP9,PF4,PPBP,VWF,MPL,FLI1,GATA1,TUBB1 |
| Immune system | Natural killer cells | NCAM1,NKG7,KLRF1,KLRD1,KLRB1,KLRC1,KLRC2,FCGR3A,GZMB,PRF1,EOMES,TBX21 | Intestine | Erythroid cells | HBB,HBA1,HBA2,HBG1,HBG2,GYPA,GYPB,ALAS2,FECH,SLC4A1,EPOR,GATA1,KLF1 |
| Immune system | Innate Lymphoid Cell 1 | PTPRC,ID2,IL7R,TBX21,IFNG,NKG7,NCR1,EOMES | Intestine | Erythroblasts | HBB,HBA1,HBA2,GYPA,ALAS2,GATA1,KLF1 |
| Immune system | Innate Lymphoid Cell 2 | PTPRC,ID2,IL7R,GATA3,IL5,IL13,IL1RL1,PTGDR2,ARG1 | Pancreas | Acinar cells | PRSS1,CPA1,CPA2,CELA3A,AMY2A,PNLIP,REG1A,CTRB1,CTRB2,KLK1 |
| Immune system | Innate Lymphoid Cell 3 | PTPRC,ID2,IL7R,RORC,IL22,IL17A,KIT,CCR6,NCR1 | Pancreas | Alpha cells | GCG,ARX,TTR,IRX2,MAFB,GC |
| Immune system | Classical Monocytes | CD14,FCGR3A,CCR2,S100A8,S100A9,LYZ,VCAN,FCN1,CD33 | Pancreas | Beta cells | INS,IAPP,MAFA,PDX1,NKX6-1,SIX2,SIX3,PCSK1 |
| Immune system | Non-classical monocytes | FCGR3A,CD14,CX3CR1,CSF1R,ITGAL,LST1,MS4A7,SIGLEC10,TCF7L2 | Pancreas | Delta cells | SST,HHEX,LEPR,RBP4 |
| Immune system | Intermediate monocytes | CD14,FCGR3A,CCR2,HLA-DRA,CD86,CCR5,ITGAX,CLEC10A | Pancreas | Ductal cells | KRT19,KRT7,SOX9,CFTR,SPP1,S100P,FOXA2,HNF1B |
| Immune system | Macrophages | CD68,CD163,MRC1,CSF1R,CD14,ITGAM,ADGRE1,APOE,C1QA,C1QB,C1QC,MARCO,MSR1 | Pancreas | Epsilon cells | GHRL |
| Immune system | Myeloid Dendritic cells | ITGAX,HLA-DRA,HLA-DRB1,HLA-DQA1,CD1C,FCER1A,CLEC10A,CD83,CD86,CCR7,IRF4,IRF8,SIRPA,CLEC9A | Pancreas | Gamma (PP) cells | PPY,PAX6 |
| Immune system | Plasmacytoid Dendritic cells | IL3RA,CLEC4C,NRP1,TCF4,IRF7,IRF8,LILRA4,GZMB,SPIB,CD123 | Pancreas | Pancreatic progenitor cells | PDX1,SOX9,NKX6-1,PTF1A,NEUROG3,HNF1B,GATA6 |
| Immune system | cDC1 | PTPRC,HLA-DRA,ITGAX,CLEC9A,XCR1,BATF3,IRF8,ID2,THBD | Pancreas | Pancreatic stellate cells | ACTA2,PDGFRB,RGS5,COL1A1,COL1A2,VIM,DES,DCN,LUM |
| Immune system | cDC2 | PTPRC,HLA-DRA,ITGAX,CD1C,SIRPA,CLEC10A,FCER1A,IRF4,NOTCH2,RELB | Pancreas | Peri-islet Schwann cells | S100B,SOX10,PLP1,MPZ,CDH19 |
| Immune system | Neutrophils | FCGR3B,CSF3R,MPO,ELANE,PRTN3,S100A8,S100A9,ITGAM,CEACAM8,CXCR1,CXCR2 | Pancreas | Endothelial cells | PECAM1,CDH5,VWF,ENG,PLVAP,KDR,FLT1 |
| Immune system | Eosinophils | CCR3,IL5RA,SIGLEC8,EPX,RNASE2,RNASE3,PRG2,CLC,HPGDS,PTGDR2 | Pancreas | Mesenchymal cells | VIM,COL1A1,COL1A2,PDGFRA,PDGFRB,DCN,LUM |
| Immune system | Basophils | FCER1A,CD200R3,ENPP3,IL3RA,CD33,TPSAB1,TPSB2,HDC,GATA2 | Pancreas | Mast cells | KIT,FCER1A,CPA3,TPSAB1,TPSB2,MS4A2 |
| Immune system | Mast cells | KIT,FCER1A,CPA3,TPSAB1,TPSB2,MS4A2,ENPP3,CD63,CD9 | Pancreas | Immune system cells | PTPRC,CD3E,CD19,CD68,NCAM1,ITGAX,KIT |
| Immune system | Granulocytes | MPO,ELANE,PRTN3,AZU1,S100A8,S100A9,CEACAM8,FCGR3B,CSF3R | Pancreas | Cancer stem cells | EPCAM,PROM1,CD44,ALDH1A1,CD24,CXCR4,SOX2,POU5F1 |
| Immune system | Platelets | ITGA2B,ITGB3,GP1BA,GP9,PPBP,PF4,SELP,TUBB1,CD9 | Kidney | Proximal tubule cells | SLC5A2,SLC5A1,LRP2,CUBN,SLC3A1,SLC7A9,SLC22A6,SLC22A8,AQP1,ALDOB,HNF4A |
| Immune system | Megakaryocyte | ITGA2B,ITGB3,GP1BA,GP9,PF4,PPBP,VWF,MPL,FLI1,GATA1,TUBB1 | Kidney | Loop of Henle cells | UMOD,SLC12A1,KCNJ1,CLCNKB,BSND |
| Immune system | Erythroid-like and erythroid precursor cells | HBB,HBA1,HBA2,HBG1,HBG2,GYPA,GYPB,ALAS2,FECH,SLC4A1,EPOR,GATA1,KLF1 | Kidney | Distal tubule cells | SLC12A3,TRPM6,KLK1,PVALB,WNK1 |
| Immune system | HSC/MPP cells | CD34,KIT,THY1,SPINK2,AVP,PROM1,HOXA9,MEIS1,GATA2,MLLT3,CD90 | Kidney | Connecting tubule cells | CALB1,TRPV5,SLC8A1,KL,AQP2,SCNN1G |
| Immune system | Progenitor cells | CD34,KIT,FLT3,MPO,AZU1,PRTN3,ELANE,GATA1,GATA2,MYB | Kidney | Principal cells (Collecting duct system) | AQP2,AQP3,AQP4,FXYD4,HSD11B2,SCNN1A,SCNN1B,SCNN1G |
| Immune system | Endothelial | PECAM1,CDH5,VWF,ENG,ICAM1,SELE,SELP,CLDN5 | Kidney | Inner Medullary cells (Collecting duct system) | AQP2,HSD11B2,SCNN1G,SLC14A2 |
| Immune system | ISG expressing immune cells | ISG15,MX1,MX2,IFIT1,IFIT2,IFIT3,IFI6,IFI44L,OAS1,OAS2,OAS3,RSAD2,STAT1 | Kidney | α-intercalated cells (Collecting duct system) | ATP6V1G3,ATP6V0D2,SLC4A1,FOXI1,INS RR |
| Blood | Naive B cells | CD19,MS4A1,CD79A,IGHM,IGHD,CD27,SELL,IL4R,CCR7,BACH2 | Kidney | β-intercalated cells (Collecting duct system) | SLC26A4,FOXI1,INS RR |
| Blood | Memory B cells | CD19,MS4A1,CD79A,CD27,CD38,SDC1,IGHG1,IGHG2,IGHG3,IGHG4,IGHA1,IGHA2,AICDA,BANK1,PAX5 | Kidney | Podocytes | NPHS1,NPHS2,PODXL,WT1,SYNPO,VEGFA,MAFB,CD2AP |
| Blood | Plasma B cells | SDC1,CD38,XBP1,PRDM1,SLAMF7,DERL3,MZB1,IGHG1,IGHG2,IGHG3,IGHG4,IGHA1,IGHA2,IGLC2,IGKC | Kidney | Mesangial cells | PDGFRB,ACTA2,ITGA8,GATA3,MEIS1,REN |
| Blood | Naive CD8+ T cells | CD3D,CD3E,CD3G,CD8A,CD8B,CCR7,SELL,LEF1,TCF7,IL7R,CD27,CD45RA | Kidney | Juxtaglomerular cells | REN,ACTA2,GJA5,CDH11,PDE3B,PVALB |
| Blood | Naive CD4+ T cells | CD3D,CD3E,CD3G,CD4,CCR7,SELL,LEF1,TCF7,IL7R,CD27,CD45RA,FOXP1 | Kidney | Endothelial cells | PECAM1,CDH5,EMCN,KDR,FLT1,VWF |
| Blood | Memory CD8+ T cells | CD3D,CD3E,CD3G,CD8A,CD8B,CD44,IL7R,CD69,EOMES,TBX21,IFNG,GZMK,GZMB,CD45RO | Kidney | Stromal cells | PDGFRA,COL1A1,COL1A2,DCN,LUM,MEIS1,FOXD1 |
| Blood | Memory CD4+ T cells | CD3D,CD3E,CD3G,CD4,CD44,IL7R,CD69,ICOS,CCR5,CCR6,CXCR3,TBX21,GATA3,RORC,FOXP3,CD45RO | Kidney | Renal interstitium (Mesenchymal cells) | PDGFRA,COL1A1,DCN,LUM,FOXD1,CLDN1,VIM |
| Blood | Effector CD8+ T cells | CD3D,CD3E,CD3G,CD8A,CD8B,GZMB,PRF1,IFNG,KLRG1,CX3CR1,TBX21,EOMES,CCL5,CD27,SELL | Kidney | Cap mesenchyme cells (Mesenchymal cells) | SIX2,CITED1,EYA1,GDNF,OSR1 |
| Blood | Effector CD4+ T cells | CD3D,CD3E,CD3G,CD4,IFNG,TNF,IL2,IL4,IL17A,IL21,ICOS,CD40LG,TBX21,GATA3,RORC,BCL6,PRDM1 | Kidney | Ureteric Bud cells | RET,WNT11,GFRA1,HOXB7,CALB1,KRT19,EPCAM |
| Blood | T regulatory cells | FOXP3,IL2RA,CTLA4,IKZF2,TNFRSF18,CD4,PTPRC,LAG3 | Kidney | Renal Vesicle cells | LHX1,PAX8,WT1,CDH6 |
| Blood | γδ-T cells | CD3D,CD3E,CD3G,TRDC,TRGC1,TRGC2,NKG7,KLRB1,KLRG1,CCL5,CD160 | Kidney | Hematopoietic cells | PTPRC,CD34,KIT,GYPA,HBB |
| Blood | MAIT cells | PTPRC,CD3D,SLC4A10,KLRB1,TRAV1-2,IL18R1,CCR6,RORC,ZBTB16 | Kidney | Immune cells | PTPRC,CD68,CD3E,CD19,ITGAM,FCGR3A,NCAM1 |
| Blood | Invariant NKT cells | PTPRC,CD3D,ZBTB16,TRAV10,TRBV25,SLAMF1,SLAMF6,CD4,CD8A | Kidney | Cancer stem cells | PROM1,CD44,ALDH1A1,EPCAM,SOX2,POU5F1,NANOG,ABCG2,CD133(PROM1) |
| Blood | CD8+ NKT-like cells | CD3D,CD8A,CD8B,NCAM1,NKG7,KLRB1,KLRD1,GZMB,PRF1,TBX21,EOMES,SLAMF7 | Brain | Astrocytes | GFAP,AQP4,SLC1A2,SLC1A3,S100B,ALDH1L1,ALDOC,SOX9,FGFR3 |
| Blood | CD4+ NKT-like cells | CD3D,CD4,NCAM1,NKG7,KLRB1,GZMK,TBX21,EOMES,SLAMF6 | Brain | Oligodendrocytes | MBP,PLP1,MOG,MAG,CNP,OLIG1,OLIG2,SOX10,NKX6-2,KLK6 |
| Blood | Natural killer cells | NCAM1,NKG7,KLRF1,KLRD1,KLRB1,KLRC1,KLRC2,FCGR3A,GZMB,PRF1,EOMES,TBX21 | Brain | Oligodendrocyte precursor cells | PDGFRA,CSPG4,SOX10,OLIG1,OLIG2,ASCL1,NKX2-2 |
| Blood | Eosinophils | CCR3,IL5RA,SIGLEC8,EPX,RNASE2,RNASE3,PRG2,CLC,HPGDS,PTGDR2 | Brain | Microglial cells | AIF1,CSF1R,CX3CR1,P2RY12,TMEM119,CD68,HEXB,C1QA,SPI1,IRF8 |
| Blood | Neutrophils | FCGR3B,CSF3R,MPO,ELANE,PRTN3,S100A8,S100A9,ITGAM,CEACAM8,CXCR1,CXCR2,SELL | Brain | Glutamatergic neurons | SLC17A6,SLC17A7,SLC17A8,TBR1,SATB2,CUX1,CUX2,BCL11B,FEZF2,FOXP2 |
| Blood | Basophils | FCER1A,CD200R3,ENPP3,IL3RA,CD33,TPSAB1,TPSB2,HDC,GATA2 | Brain | GABAergic neurons | GAD1,GAD2,SLC32A1,DLX1,DLX2,DLX5,ARX,LHX6,SST,NPY,VIP,CALB1,CALB2,PVALB |
| Blood | Classical Monocytes | CD14,FCGR3A,CCR2,S100A8,S100A9,LYZ,VCAN,FCN1,CD33,SELL | Brain | Dopaminergic neurons | TH,SLC6A3,SLC18A2,DDC,NR4A2,EN1,LMX1A,LMX1B,PITX3 |
| Blood | Non-classical monocytes | FCGR3A,CD14,CX3CR1,CSF1R,ITGAL,LST1,MS4A7,SIGLEC10,TCF7L2,ITGAM | Brain | Serotonergic neurons | TPH2,SLC6A4,FEV,GATA3,LMX1B,PET1(FEV) |
| Blood | Intermediate monocytes | CD14,FCGR3A,CCR2,HLA-DRA,CD86,CCR5,ITGAX,CLEC10A | Brain | Cholinergic neurons | CHAT,SLC5A7,SLC18A3,ACHE,NGFR |
| Blood | cDC1 | PTPRC,HLA-DRA,ITGAX,CLEC9A,XCR1,BATF3,IRF8,ID2,THBD | Brain | Mature neurons | RBFOX3,TUBB3,MAP2,SYP,SYN1,SNAP25,NEFL,NEFM,NEFH |
| Blood | cDC2 | PTPRC,HLA-DRA,ITGAX,CD1C,SIRPA,CLEC10A,FCER1A,IRF4,NOTCH2,RELB | Brain | Immature neurons | DCX,TUBB3,STMN2,NEUROD1,NEUROD2,NEUROG1,NEUROG2,ELAVL4 |
| Blood | Plasmacytoid Dendritic cells | IL3RA,CLEC4C,NRP1,TCF4,IRF7,IRF8,LILRA4,GZMB,SPIB,CD123,HLA-DRA | Brain | Neuroblasts | DCX,TUBB3,STMN2,NEUROD1,TAGLN3,ASCL1 |
| Blood | HSC/MPP cells | CD34,KIT,THY1,SPINK2,AVP,PROM1,HOXA9,MEIS1,GATA2,MLLT3,CD90 | Brain | Neural Progenitor cells | NES,SOX2,PAX6,OLIG2,ASCL1,MSI1,FABP7,HES1,HES5 |
| Blood | Progenitor cells | CD34,KIT,FLT3,MPO,AZU1,PRTN3,ELANE,GATA1,GATA2,MYB | Brain | Neural stem cells | NES,SOX2,PAX6,PROM1,GFAP,ID1,MSI1,FABP7,HOPX |
| Blood | Erythrocytes | HBB,HBA1,HBA2,GYPA,GYPB,SLC4A1,ALAS2,FECH,EPB41,SPTA1,CA1 | Brain | Neuroepithelial cells | SOX2,PAX6,NES,HES1,HES5,CDH2 |
| Blood | Reticulocytes | HBB,HBA1,HBA2,GYPA,TFRC,ALAS2,RPS27,RPS19,RPL3,FTL,SLC4A1 | Brain | Radial glial cells | GFAP,VIM,NES,SOX2,PAX6,FABP7,SLC1A3,HES1,HES5 |
| Blood | Platelets | ITGA2B,ITGB3,GP1BA,GP9,PPBP,PF4,SELP,TUBB1,CD9,CLU | Brain | Ependymal Cells | FOXJ1,RSPH1,PIFO,CD24,GFAP,S100B,VIM,TMEM212 |
| Liver | Hepatocytes | ALB,ASGR1,CPS1,HNF4A | Brain | Choroid Plexus Epithelial Cells | TTR,AQP1,KRT8,KRT18,OTX2,ENPP2,CLIC6,FOLR1,CDH1 |
| Liver | Cholangiocytes | KRT19,KRT7,EPCAM,SOX9,CFTR,CLDN4,AQP1,SLC10A2,NR1H4 | Brain | Tanycytes | RAX,COL25A1,DIO2,SLC16A2,GFAP,VIM,PPP1R1B |
| Liver | Hepatoblasts | AFP,EPCAM,KRT19,KRT8,DLK1,GPC3,ALB | Brain | Myelinating Schwann cells | MPZ,MBP,PMP22,MAG,PLP1,SOX10,EGR2,POU3F1 |
| Liver | Hepatic Stellate Cells | LRAT,RBP1,RELN,PDGFRB,DES,NGFR,GFAP,ACTA2,COL1A1 | Brain | Non myelinating Schwann cells | S100B,SOX10,NCAM1,CDH19,GFAP,NGFR |
| Liver | Kupffer cells | CD68,MARCO,CD163,CSF1R,C1QA,C1QB,CLEC4F,TIMD4,VSIG4,ITGAM | Brain | Schwann precursor cells | SOX10,FABP7,MPZ,PLP1,ERBB3 |
| Liver | Liver progenitor cell | EPCAM,KRT19,SOX9,PROM1,CD24,AFP,LGR5 | Brain | Endothelial cells | PECAM1,CDH5,CLDN5,VWF,SLC2A1,ABCB1,ICAM1,PLVAP |
| Liver | Pit cells | PTPRC,CD3D,NCAM1,NKG7,KLRF1,FCGR3A,CXCR6,ITGA1,PRF1,GZMB | Brain | Immune system cells | PTPRC,AIF1,CD68,CSF1R,CX3CR1,P2RY12,TMEM119,CD3E,MS4A1 |
| Liver | Endothelial cell | PECAM1,CDH5,STAB1,STAB2,LYVE1,VWF,KDR,ICAM1,CLEC4G,CLEC4M | Brain | Cancer cells | GFAP,OLIG2,EGFR,PDGFRA,IDH1,IDH2,ATRX,MKI67,SOX2,NES,MYC,OTX2 |
| Liver | Hematopoietic cell | PTPRC,CD34,KIT,RUNX1,SPN | Brain | Cancer stem cells | PROM1,CD44,NES,SOX2,OLIG2,L1CAM,ALDH1A1,MSI1 |
| Liver | Hematopoeitic precursor | CD34,CD38,CD133,CD117,CD90,CD48,CD150 | Eye | Rod photoreceptor cells | RHO,SAG,PDE6B,PDE6A,GNGT1,ROM1,PRPH2,NRL,NR2E3,RCVRN |
| Liver | Fibroblasts | PDGFRA,FBLN2,DCN,LUM,THY1,CD34,COL1A1,COL1A2,VIM | Eye | Cone photoreceptor cells | OPN1SW,OPN1MW,OPN1LW,ARR3,PDE6C,PDE6H,GNGT2,RXRG,THRB,RCVRN |
| Liver | Mesenchymal stem cell | ENG,NT5E,THY1,PDGFRA,VIM,CD44,NES | Eye | Retinal ganglion cells | POU4F1,POU4F2,THY1,NEFL,NEFM,SNCG,GAP43,TUBB3,RBPMS,SLC17A6 |
| Liver | Erythrocytes | HBB,HBA1,HBA2,GYPA,GYPB,SLC4A1,ALAS2,FECH,EPB41,SPTA1,CA1 | Eye | Rod bipolar cells | PRKCA,CABP5,TRPM1,GRIK1,LRIT3,PCP2 |
| Liver | Mast cells | KIT,FCER1A,CPA3,TPSAB1,TPSB2,MS4A2 | Eye | Cone bipolar cells | PRKCA,CABP5,GRIK1,ISL1,GRM6,VSX1,VSX2,NKX2-2 |
| Liver | Neutrophils | FCGR3B,CSF3R,MPO,ELANE,PRTN3,S100A8,S100A9,ITGAM,CEACAM8,CXCR1,CXCR2,SELL | Eye | Horizontal cells | CALB1,LHX1,PROX1,PCP4,ONECUT1,ONECUT2 |
| Liver | Naive B cells | CD19,MS4A1,CD79A,IGHM,IGHD,CD27,SELL,IL4R,CCR7,BACH2 | Eye | GABAergic amacrine cells | GAD1,GAD2,SLC32A1,PAX6,CALB1,CALB2,NPY,SOM,VIP |
| Liver | Memory B cells | CD19,MS4A1,CD79A,CD27,CD38,SDC1,IGHG1,IGHG2,IGHG3,IGHG4,IGHA1,IGHA2,AICDA,BANK1,PAX5 | Eye | Glycinergic amacrine cells | SLC6A9,PAX6,SLC32A1,EBF3 |
| Liver | Plasma B cells | SDC1,CD38,XBP1,PRDM1,SLAMF7,DERL3,MZB1,IGHG1,IGHG2,IGHG3,IGHG4,IGHA1,IGHA2,IGLC2,IGKC | Eye | Cholinergic GABAergic amacrine cells | CHAT,SLC5A7,SLC18A3,GAD1,GAD2,SLC32A1 |
| Liver | CD8+ NKT-like cells | CD3D,CD8A,CD8B,NCAM1,NKG7,KLRB1,KLRD1,GZMB,PRF1,TBX21,EOMES,SLAMF7 | Eye | Tachykinin GABAergic amacrine cells | TAC1,GAD1,GAD2,SLC32A1 |
| Liver | CD4+ NKT-like cells | CD3D,CD4,NCAM1,NKG7,KLRB1,GZMK,TBX21,EOMES,SLAMF6 | Eye | Starburst amacrine cells | CHAT,SLC18A3,SLC5A7,FEZF1,GAD1,GAD2,SLC32A1 |
| Liver | Naive CD8+ T cells | CD3D,CD3E,CD3G,CD8A,CD8B,CCR7,SELL,LEF1,TCF7,IL7R,CD27,CD45RA | Eye | Muller cells | RLBP1,GLUL,SLC1A3,AQP4,VIM,CD44,SOX2,PAX6 |
| Liver | Naive CD4+ T cells | CD3D,CD3E,CD3G,CD4,CCR7,SELL,LEF1,TCF7,IL7R,CD27,CD45RA,FOXP1 | Eye | Retinal pigment epithelial cells | RPE65,BEST1,RLBP1,PMEL,TYR,DCT,MITF,KRT8,KRT18,TRPM1 |
| Liver | Memory CD8+ T cells | CD3D,CD3E,CD3G,CD8A,CD8B,CD44,IL7R,CD69,EOMES,TBX21,IFNG,GZMK,GZMB,CD45RO | Eye | Retinal progenitor cells | PAX6,SOX2,VSX2,LHX2,NES,CCND1,ASCL1,OLIG2 |

| Tissue | Cell Type | Markers | Tissue | Cell Type | Markers |
|---|---|---|---|---|---|
| Liver | Memory CD4+ T cells | CD3D,CD3E,CD3G,CD4,CD44,IL7R,CD69,ICOS,CCR5,CCR6,CXCR3,TBX21,GATA3,RORC,FOXP3,CD45RO | Eye | Cornea epithelial cells | KRT12,KRT3,KRT5,KRT14,TP63,PAX6,EPCAM |
| Liver | Effector CD8+ T cells | CD3D,CD3E,CD3G,CD8A,CD8B,GZMB,PRF1,IFNG,KLRG1,CX3CR1,TBX21,EOMES,CCL5,CD27,SELL | Eye | Corneal Endothelial Cells | SLC4A11,AQP1,PITX2,COL8A2,CDH2,ATP1A1,TJP1 |
| Liver | Effector CD4+ T cells | CD3D,CD3E,CD3G,CD4,IFNG,TNF,IL2,IL4,IL17A,IL21,ICOS,CD40LG,TBX21,GATA3,RORC,BCL6,PRDM1 | Eye | Lens Epithelial Cells | CRYAA,CRYAB,PAX6,FOXE3,KRT19,HSF4,CDH1,LIMS1 |
| Liver | T regulatory cells | FOXP3,IL2RA,CTLA4,IKZF2,TNFRSF18,CD4,PTPRC,LAG3 | Eye | Astrocytes | GFAP,AQP4,SLC1A2,S100B,ALDH1L1,ALDOC |
| Liver | γδ-T cells | CD3D,CD3E,CD3G,TRDC,TRGC1,TRGC2,NKG7,KLRB1,KLRG1,CCL5,CD160 | Eye | Microglial cells | AIF1,CD68,CSF1R,CX3CR1,P2RY12,TMEM119,HEXB,C1QA |
| Liver | Classical Monocytes | CD14,FCGR3A,CCR2,S100A8,S100A9,LYZ,VCAN,FCN1,CD33 | Eye | Endothelial cells | PECAM1,CDH5,VWF,ICAM1,CLDN5,PLVAP |
| Liver | Non-classical monocytes | FCGR3A,CD14,CX3CR1,CSF1R,ITGAL,LST1,MS4A7,SIGLEC10,TCF7L2 | Eye | Pericytes | PDGFRB,RGS5,ACTA2,MCAM,CSPG4,KCNJ8,ABCC9,CD146 |
| Liver | Intermediate monocytes | CD14,FCGR3A,CCR2,HLA-DRA,CD86,CCR5,ITGAX,CLEC10A | Eye | Fibroblasts | COL1A1,COL1A2,VIM,DCN,LUM,PDGFRA,FN1 |
| Liver | Myeloid Dendritic cells | ITGAX,HLA-DRA,HLA-DRB1,HLA-DQA1,CD1C,FCER1A,CLEC10A,CD83,CD86,CCR7,IRF4,IRF8,SIRPA,CLEC9A | Eye | Immune cells | PTPRC,AIF1,CD68,CD3E,CD19,ITGAM |
| Liver | Plasmacytoid Dendritic cells | IL3RA,CLEC4C,NRP1,TCF4,IRF7,IRF8,LILRA4,GZMB,SPIB,CD123 | Eye | Synapses | SYP,SYN1,SNAP25,STX1A,SYT1,CADM1,NLGN1,NRXN1 |
| Lung | Cancer stem cells | EPCAM,PROM1,CD44,ALDH1A1,CD24,THY1,NANOG,SOX2 | Eye | Extracellular matrix cells | COL1A1,COL4A1,FN1,LAMB1,VCAN,DCN,LUM,ELN |
| Lung | Pulmonary alveolar type I cells | AGER,PDPN,CAV1,ICAM1,CLIC5,HOPX,AQP5 | Eye | Cancer cells | PMEL,MLANA,SOX10,MITF,TYR,MKI67,RB1,CRX,ARR3 |
| Lung | Pulmonary alveolar type II cells | SFTPC,SFTPB,SFTPA1,SFTPA2,SFTPD,LAMP3,ABCA3,ETV5,SOX9,NAPSA | Heart | Cardiomyocytes | TNNT2,TNNC1,TNNI3,MYH6,MYH7,ACTC1,NPPA,NPPB,MYL2,MYL7,NKX2-5,GATA4,TBX5 |
| Lung | Clara cells | SCGB1A1,SCGB3A2,CYP2F1,UPK3A,CEACAM6 | Heart | Cardiac Conduction System Cells | HCN4,TBX3,TBX5,SHOX2,ISL1,SCN5A,CACNA1H,GJA5,KCND2,NKX2-5 |
| Lung | Basal cells | KRT5,KRT14,KRT15,TP63,NGFR,SOX2,NOTCH1 | Heart | Purkinje Fibers | GJA5,SCN5A,KCND2,ID2,RYR2,MYL2,ATP2A2,TNNT2,ACTC1 |
| Lung | Ciliated cells | FOXJ1,PIFO,DYNC2LI1,DNAH5,CCDC39,RSPH1,TP73 | Heart | Endocardial cells | NPR3,CDH5,PECAM1,NFATC1,HEY1,MSX1 |
| Lung | Lung multiciliated epithelial cell | FOXJ1,TPPP3,CADPS,PIFO,DNAH5 | Heart | Vascular endothelial cells | PECAM1,CDH5,VWF,ICAM1,ENG,KDR,FLT1 |
| Lung | Ionocytes | FOXI1,CFTR,ASCL3,ATP6V0D2,STAP1 | Heart | Lymphatic endothelial cells | PROX1,LYVE1,PDPN,FLT4,CCL21,PECAM1,CDH5 |
| Lung | Airway goblet cells | MUC5AC,MUC5B,SPDEF,FOXA3,TFF1,TFF2,TFF3,REG4 | Heart | Stromal cells | PDGFRA,PDGFRB,DCN,LUM,COL1A1,COL1A2,VIM,TCF21,POSTN |
| Lung | Goblet / Secretory cell | SCGB1A1,MUC5AC,MUC5B,SCGB3A1,SCGB3A2,UPK3A | Heart | Smooth muscle cells | ACTA2,TAGLN,MYH11,CNN1,DES,CALD1 |
| Lung | Bronchus Serous cell | LPO,BPIFB1,LTF,PRR4,SLPI,BPIFA1 | Heart | Epicardial fat cells | ADIPOQ,PLIN1,LEP,FABP4,LIPE,CEBPA,PPARG |
| Lung | Epithelial cells | EPCAM,KRT8,KRT18,CDH1 | Heart | Schwann cells | S100B,SOX10,PLP1,MPZ,GFAP |
| Lung | Pulmonary Neuroendocrine Cells | ASCL1,CHGA,CHGB,SYP,GRP,CALCA,UCHL1,INSM1,EPCAM | Heart | Visceral neurons | TH,CHAT,NOS1,TUBB3,ELAVL4,PHOX2B,NGF |
| Lung | Mesothelial cells | MSLN,WT1,UPK3B,CALB2,LRRN4 | Heart | Myeloid cells | PTPRC,CD68,MRC1,CD163,ITGAM,CSF1R,ADGRE1,CCR2 |
| Lung | Mesothelial cells | MSLN,WT1,UPK3B,CALB2,KRT8,KRT18,CDH1 | Heart | Lymphoid cells | PTPRC,CD3E,CD8A,CD4,CD19,MS4A1,NCAM1,CCR7,IL7R |
| Lung | Endothelial cell | PECAM1,CDH5,VWF,ICAM1,EMCN,ACKR1,PLVAP,CA4 | Heart | Megakaryocytes | ITGA2B,ITGB3,GP1BA,PF4,PPBP |
| Lung | Capillary endothelial cells | EDNRB,CA4,CD36,SGK1,RGCC | Heart | Erythroblasts | HBB,HBA1,HBA2,GYPA,ALAS2,GATA1,KLF1 |
| Lung | Capillary endothelial cell | PECAM1,CA4,RGCC,SGK1,PRX | Heart | SATB2_LRRC7 positive cells | SATB2,LRRC7 |
| Lung | Bronchial endothelial cells | VWF,ACKR1,SELE,BMPER | Heart | ELF3_AGBL2 positive cells | ELF3,AGBL2 |
| Lung | Fibroblasts | COL1A1,COL1A2,COL3A1,PDGFRA,PDGFRB,DCN,LUM,VIM,FN1,ACTA2 | Heart | CLC_IL5RA positive cells | CLC,IL5RA |
| Lung | Adventitial fibroblasts | PI16,MFAP5,SCARA5,DCN,LUM | Adrenal | Adrenocortical cells | CYP11B1,CYP11B2,CYP17A1,CYP21A2,HSD3B2,MC2R,NR5A1,STAR |
| Lung | Pericytes | COX4I2,PDGFRB,RGS5,CSPG4,KCNJ8 | Adrenal | Chromaffin cells | CHGA,CHGB,TH,DBH,PNMT,SYP,SLC18A1 |
| Lung | Vascular associated smooth muscle cell | ACTA2,MYH11,NOTCH3,CNN1,RGS5 | Adrenal | Sympathoblasts | TH,PHOX2B,DBH,ASCL1,SOX10,CHGA |
| Lung | Bronchial smooth muscle cell | ACTA2,MYH11,HHIP,DES,CALD1 | Adrenal | Schwann cells | S100B,SOX10,PLP1,MPZ,CDH19 |
| Lung | Alveolar macrophages | CD68,MRC1,MARCO,MSR1,FABP4,PPARG,SIGLEC1,SIGLECF | Adrenal | Stromal cells | COL1A1,COL1A2,VIM,DCN,PDGFRA,PDGFRB |
| Lung | Macrophages | CD68,CD163,MRC1,CSF1R,CD14,ITGAM,ADGRE1,APOE,C1QA,C1QB,C1QC,MARCO,MSR1 | Adrenal | Vascular endothelial cells | PECAM1,CDH5,VWF,ENG,PLVAP |
| Lung | Classical Monocytes | CD14,FCGR3A,CCR2,S100A8,S100A9,LYZ,VCAN,FCN1,CD33 | Adrenal | Myeloid cells | PTPRC,CD68,ITGAM,ADGRE1,CSF1R,CD14,FCGR3A |
| Lung | Non-classical monocytes | FCGR3A,CD14,CX3CR1,CSF1R,ITGAL,LST1,MS4A7,SIGLEC10,TCF7L2 | Adrenal | Lymphoid cells | PTPRC,CD3E,CD8A,CD4,CD19,MS4A1,NCAM1 |
| Lung | Intermediate monocytes | CD14,FCGR3A,CCR2,HLA-DRA,CD86,CCR5,ITGAX,CLEC10A | Adrenal | Megakaryocytes | ITGA2B,ITGB3,GP1BA,PF4,PPBP |
| Lung | Myeloid Dendritic cells | ITGAX,HLA-DRA,HLA-DRB1,HLA-DQA1,CD1C,FCER1A,CLEC10A,CD83,CD86,CCR7,IRF4,IRF8,SIRPA,CLEC9A | Adrenal | Erythroblasts | HBB,HBA1,HBA2,GYPA,ALAS2,GATA1,KLF1 |
| Lung | Plasmacytoid Dendritic cells | IL3RA,CLEC4C,NRP1,TCF4,IRF7,IRF8,LILRA4,GZMB,SPIB,CD123 | Adrenal | CSH1_CSH2 positive cells | CSH1,CSH2 |
| Lung | Granulocytes | MPO,ELANE,PRTN3,AZU1,S100A8,S100A9,CEACAM8,FCGR3B,CSF3R | Adrenal | SLC26A4_PAEP positive cells | SLC26A4,PAEP |
| Lung | Neutrophils | FCGR3B,CSF3R,MPO,ELANE,PRTN3,S100A8,S100A9,ITGAM,CEACAM8,CXCR1,CXCR2 | Muscle | Skeletal muscle cells | MYH1,MYH2,MYH4,MYH7,ACTN2,ACTN3,TNNT1,TNNT3,CKM,DES,MYOD1,MYOG |
| Lung | Eosinophils | CCR3,IL5RA,SIGLEC8,EPX,RNASE2,RNASE3,PRG2,CLC,HPGDS,PTGDR2 | Muscle | Satellite cells | PAX7,MYF5,CD34,CAV1,VCAM1,NCAM1,ITGA7 |
| Lung | Basophils | FCER1A,CD200R3,ENPP3,IL3RA,CD33,TPSAB1,TPSB2,HDC,GATA2 | Muscle | Fibro-adipogenic Progenitors | PDGFRA,LY6A,VIM,COL1A1,DCN,CD34,PPARG,CEBPA |
| Lung | Mast cells | KIT,FCER1A,CPA3,TPSAB1,TPSB2,MS4A2,ENPP3,CD63,CD9 | Muscle | Smooth muscle cells | ACTA2,TAGLN,MYH11,CNN1,DES |
| Lung | Naive B cells | CD19,MS4A1,CD79A,IGHM,IGHD,CD27,SELL,IL4R,CCR7,BACH2 | Muscle | Stromal cells | PDGFRA,COL1A1,VIM,DCN,LUM,TCF4,ASPN |
| Lung | Memory B cells | CD19,MS4A1,CD79A,CD27,CD38,SDC1,IGHG1,IGHG2,IGHG3,IGHG4,IGHA1,IGHA2,AICDA,BANK1,PAX5 | Muscle | Vascular endothelial cells | PECAM1,CDH5,VWF,ENG,KDR,ICAM1 |
| Lung | Plasma B cells | SDC1,CD38,XBP1,PRDM1,SLAMF7,DERL3,MZB1,IGHG1,IGHG2,IGHG3,IGHG4,IGHA1,IGHA2,IGLC2,IGKC | Muscle | Lymphatic endothelial cells | PROX1,LYVE1,PDPN,FLT4,PECAM1,CDH5 |
| Lung | Naive CD8+ T cells | CD3D,CD3E,CD3G,CD8A,CD8B,CCR7,SELL,LEF1,TCF7,IL7R,CD27,CD45RA | Muscle | Schwann cells | S100B,SOX10,PLP1,MPZ,GFAP,NGFR |
| Lung | Naive CD4+ T cells | CD3D,CD3E,CD3G,CD4,CCR7,SELL,LEF1,TCF7,IL7R,CD27,CD45RA,FOXP1 | Muscle | Myeloid cells | PTPRC,CD68,MRC1,ITGAM,ADGRE1,CSF1R |
| Lung | Memory CD8+ T cells | CD3D,CD3E,CD3G,CD8A,CD8B,CD44,IL7R,CD69,EOMES,TBX21,IFNG,GZMK,GZMB,CD45RO | Muscle | Lymphoid cells | PTPRC,CD3E,CD4,CD8A,NCAM1 |
| Lung | Memory CD4+ T cells | CD3D,CD3E,CD3G,CD4,CD44,IL7R,CD69,ICOS,CCR5,CCR6,CXCR3,TBX21,GATA3,RORC,FOXP3,CD45RO | Muscle | Megakaryocytes | ITGA2B,ITGB3,GP1BA,PF4,PPBP |
| Lung | Effector CD8+ T cells | CD3D,CD3E,CD3G,CD8A,CD8B,GZMB,PRF1,IFNG,KLRG1,CX3CR1,TBX21,EOMES,CCL5,CD27,SELL | Muscle | Erythroblasts | HBB,HBA1,HBA2,GYPA,ALAS2,GATA1,KLF1 |
| Lung | Effector CD4+ T cells | CD3D,CD3E,CD3G,CD4,IFNG,TNF,IL2,IL4,IL17A,IL21,ICOS,CD40LG,TBX21,GATA3,RORC,BCL6,PRDM1 | Stomach | Parietal and chief cells | ATP4A,ATP4B,GIF,PGA3,PGA4,PGC,LIPF,BHLHA15 |
| Lung | gamma delta-T cells | CD3D,CD3E,CD3G,TRDC,TRGC1,TRGC2,NKG7,KLRB1,KLRG1,CCL5,CD160 | Stomach | Neuroendocrine cells | CHGA,CHGB,SYP,NCAM1,SST,GAST,GHRL,TPH1,INS L1 |
| Lung | CD8+ NKT-like cells | CD3D,CD8A,CD8B,NCAM1,NKG7,KLRB1,KLRD1,GZMB,PRF1,TBX21,EOMES,SLAMF7 | Stomach | Goblet cells | MUC2,TFF3,FCGBP,SPDEF |
| Lung | CD4+ NKT-like cells | CD3D,CD4,NCAM1,NKG7,KLRB1,GZMK,TBX21,EOMES,SLAMF6 | Stomach | Ciliated epithelial cells | FOXJ1,PIFO,KRT8,KRT18,EPCAM |
| Lung | Natural killer cells | NCAM1,NKG7,KLRF1,KLRD1,KLRB1,KLRC1,KLRC2,FCGR3A,GZMB,PRF1,EOMES,TBX21 | Stomach | Squamous epithelial cells | KRT5,KRT14,TP63 |
| Lung | Platelets | ITGA2B,ITGB3,GP1BA,GP9,PPBP,PF4,SELP,TUBB1,CD9 | Stomach | Stromal cells | COL1A1,COL1A2,VIM,DCN,LUM,PDGFRA,PDGFRB,POSTN |
| Lung | Megakaryocyte | ITGA2B,ITGB3,GP1BA,GP9,PF4,PPBP,VWF,MPL,FLI1,GATA1,TUBB1 | Stomach | Vascular endothelial cells | PECAM1,CDH5,VWF,ENG,PLVAP |
| Lung | Erythroid cells | HBB,HBA1,HBA2,HBG1,HBG2,GYPA,GYPB,ALAS2,FECH,SLC4A1,EPOR,GATA1,KLF1 | Stomach | Lymphatic endothelial cells | PROX1,LYVE1,PDPN,FLT4 |
| Lung | Progenitor cells | CD34,KIT,FLT3,MPO,AZU1,PRTN3,ELANE,GATA1,GATA2,MYB | Stomach | Mesothelial cells | MSLN,WT1,UPK3B,CALB2 |
| Intestine | Cancer stem cells | EPCAM,PROM1,CD44,ALDH1A1,SOX2,POU5F1,NANOG,KRT5,TP63 | Stomach | ENS neurons | TUBB3,ELAVL4,PHOX2B,CHAT,NOS1,VIP,GAL |
| Intestine | Enterocytes | FABP2,ALPI,VIL1,SI,SLC26A3,ANPEP,KRT20,FABP1,SLC5A1,ACE2 | Stomach | ENS glia | S100B,SOX10,PLP1,FABP7,GFAP |
| Intestine | Intestinal epithelial cells | EPCAM,CDH1,KRT8,KRT18,VIL1 | Stomach | Myeloid cells | PTPRC,CD68,MRC1,ITGAM,ADGRE1,CD14,KIT,FCER1A |
| Intestine | Intestinal Stem Cells | LGR5,OLFM4,ASCL2,SOX9,SMOC2,RGMB,BMI1,TERT,EPCAM | Stomach | Lymphoid cells | PTPRC,CD3E,CD4,CD8A,CD19,MS4A1,NCAM1,ITGAE |
| Intestine | Transit amplifying cells | MKI67,TOP2A,PCNA,OLFM4,EPHB2,SOX9,STMN1,MCM5,CENPF | Stomach | Erythroblasts | HBB,HBA1,HBA2,GYPA,ALAS2 |
| Intestine | Crypt cells | LGR5,ASCL2,OLFM4,SOX9,SMOC2,EPHB2,CD44,AXIN2,RGMB | Stomach | MUC13_DMBT1 positive cells | MUC13,DMBT1 |
| Intestine | Goblet cells | MUC2,TFF3,FCGBP,ZG16,CLCA1,AGR2,SPDEF | Stomach | PDE1C_ACSM3 positive cells | PDE1C,ACSM3 |
| Intestine | Paneth Cells | LYZ,DEFA5,DEFA6,MMP7,PLA2G2A,REG3A,ITLN2 | Spleen | Lymphoid cells | PTPRC,CD3E,CD4,CD8A,CD19,MS4A1,CR2,NCAM1,IGHM,IGHD,AICDA,FOXP3 |
| Intestine | Tuft Cells | POU2F3,TRPM5,IL25,AVIL,DCLK1,GFI1B,HCK,EPCAM | Spleen | Myeloid cells | PTPRC,CD68,CD163,MRC1,ITGAM,ADGRE1,CD14,ITGAX,CLEC4C,FCER1A |
| Intestine | Tuft cells | DCLK1,POU2F3,TRPM5,SH2D6,BMX,AVIL,LRMP,GFI1B,HCK,IL25 | Spleen | Stromal cells | PDGFRA,PDGFRB,COL1A1,VIM,DCN,LUM,CCL19,CCL21,MADCAM1,LTBR,CR2 |
| Intestine | M cells | SPIB,GP2,TNFRSF11A,CCL9,CKLF,MARCKSL1,ANXA5,EPCAM | Spleen | Vascular endothelial cells | PECAM1,CDH5,VWF,ENG,ACKR1,SELP |
| Intestine | Enteroendocrine cells | CHGA,CHGB,SYP,NEUROD1,TPH1,GCG,PYY,CCK,SCT,GIP,SST | Spleen | Mesothelial cells | MSLN,WT1,UPK3B,CALB2 |
| Intestine | Chromaffin cells | CHGA,TH,TPH1,DDC | Spleen | Megakaryocytes | ITGA2B,ITGB3,GP1BA,PF4,PPBP,MPL |
| Intestine | Enteroglial cells | S100B,GFAP,SOX10,PLP1,CDH19,FOXD3,GPM6B,CRYAB,ERBB3 | Spleen | Erythroblasts | HBB,HBA1,HBA2,GYPA,ALAS2,GATA1,KLF1 |
| Intestine | ENS glia | S100B,SOX10,PLP1,FABP7,GFAP,BFABP(FABP7) | Spleen | AFP_ALB positive cells | AFP,ALB |
| Intestine | Fibroblasts | COL1A1,COL1A2,COL3A1,PDGFRA,DCN,LUM,FN1,ACTA2 | Spleen | STC2_TLX1 positive cells | STC2,TLX1 |
| Intestine | Myofibroblast cells | COL1A1,POSTN,TIMP1,ACTA2,TAGLN,FN1,CNN1 | Thymus | Thymocytes | PTPRC,IL7R,CD3D,CD3E,CD4,CD8A,CD8B,CD1A,CD44,CD24,CD69,RAG1,RAG2,PTCRA,TRBC1,TRBC2,FOXP3,IKZF1 |
| Intestine | Smooth muscle cells | ACTA2,TAGLN,MYH11,CNN1,CALD1,SMTN | Thymus | Thymic epithelial cells | EPCAM,KRT8,KRT5,KRT14,CDH1,FOXN1,AIRE,CCL25,PSMB11,CD83,CD40,HLA-DRA |
| Intestine | Pericytes | RGS5,NOTCH3,MCAM,KCNJ8,ABCC9,HIGD1B,GJA4 | Thymus | Antigen presenting cells | HLA-DRA,HLA-DRB1,CD80,CD86,CD40,ITGAX,CD68,CLEC4C,IL3RA |
| Intestine | Vascular endothelial cells | PECAM1,CDH5,VWF,ENG,PLVAP,SELE | Thymus | Stromal cells | PDGFRA,COL1A1,VIM,DCN,LUM,PDPN |
| Intestine | Lymphatic endothelial cells | PROX1,LYVE1,PDPN,FLT4,CCL21,PECAM1,CDH5 | Thymus | Vascular endothelial cells | PECAM1,CDH5,VWF,ENG,SELE,ICAM1 |
| Intestine | Mesothelial cells | MSLN,WT1,UPK3B,CALB2,KRT8,KRT18 | Placenta | Syncytiotrophoblasts and villous cytotrophoblasts | KRT7,GJA1,CDH1,EPCAM,CGA,CGB,CYP19A1,ERVFRD-1,GCM1,TEAD4 |
| Intestine | Innate Lymphoid Cell 1 | PTPRC,ID2,IL7R,TBX21,IFNG,NKG7,NCR1,EOMES | Placenta | Extravillous trophoblasts | HLA-G,ITGA5,ITGA1,CDH1,KRT7,MMP2,MMP9,PAPPA2 |
| Intestine | Innate Lymphoid Cell 2 | PTPRC,ID2,IL7R,GATA3,IL5,IL13,IL1RL1,PTGDR2,ARG1 | Placenta | Trophoblast giant cells | HAND1,ASCL2,GCM1,KRT7,CDH1 |
| Intestine | Innate Lymphoid Cell 3 | PTPRC,ID2,IL7R,RORC,IL22,IL17A,KIT,CCR6,NCR1 | Placenta | Stromal cells | COL1A1,COL1A2,VIM,PDGFRA,PDGFRB,DCN,LUM |
| Intestine | Myeloid cells | PTPRC,CD68,MRC1,ITGAM,CSF1R,ADGRE1,CD14,FCGR3A,ITGAX,KIT | Placenta | Vascular endothelial cells | PECAM1,CDH5,VWF,ENG,KDR |
| Intestine | Macrophages | CD68,CD163,MRC1,CSF1R,CD14,ITGAM,ADGRE1,APOE,C1QA,C1QB,C1QC,MARCO,MSR1 | Placenta | Smooth muscle cells | ACTA2,TAGLN,MYH11,CNN1 |
| Intestine | Classical Monocytes | CD14,FCGR3A,CCR2,S100A8,S100A9,LYZ,VCAN,FCN1,CD33 | Placenta | Myeloid cells | PTPRC,CD68,MRC1,CD163,CSF1R,ITGAM,CD14,LYZ |
| Intestine | Non-classical monocytes | FCGR3A,CD14,CX3CR1,CSF1R,ITGAL,LST1,MS4A7,SIGLEC10,TCF7L2 | Placenta | Lymphoid cells | PTPRC,CD3E,CD8A,CD4,NCAM1,TRDC |
| Intestine | Intermediate monocytes | CD14,FCGR3A,CCR2,HLA-DRA,CD86,CCR5,ITGAX,CLEC10A | Placenta | Megakaryocytes | ITGA2B,ITGB3,GP1BA,PF4,PPBP |
| Intestine | Myeloid Dendritic cells | ITGAX,HLA-DRA,HLA-DRB1,HLA-DQA1,CD1C,FCER1A,CLEC10A,CD83,CD86,CCR7,IRF4,IRF8,SIRPA,CLEC9A | Placenta | PAEP_MECOM positive cells | PAEP,MECOM |
| Intestine | Plasmacytoid Dendritic cells | IL3RA,CLEC4C,NRP1,TCF4,IRF7,IRF8,LILRA4,GZMB,SPIB,CD123 | Placenta | AFP_ALB positive cells | AFP,ALB |
| Intestine | Neutrophils | FCGR3B,CSF3R,MPO,ELANE,PRTN3,S100A8,S100A9,ITGAM,CEACAM8,CXCR1,CXCR2 | Placenta | IGFBP1_DKK1 positive cells | IGFBP1,DKK1 |

**Supplementary Table 3. Summary statistics comparing max mean annotation accuracy of CellTypeAI:model combinations (and Google's Gemini 2.5 Pro using CellTypeAI's engineered prompt) versus conventional annotation methods (CellTypist and ScType).** n = 3. Where both groups' standard deviation (s.d.) = 0, means were directly compared (p set to 0.0 for unequal means). Where one group's s.d. = 0, a one-sample t-test was used. Where both groups' s.d. >0, a two-tailed Welch's t-test was used. P < 0.05 was considered significant. ns = not significant | * p < 0.05 | ** p < 0.01 | *** p < 0.001.

| Comparison | Dataset | t-stat | p-value | Significance |
|---|---|---|---|---|
| CellTypeAI: Qwen3:235b vs CellTypist | PBMC3K | 6.386937353 | 0.023647937 | * |
| CellTypeAI: Qwen3:235b vs ScType | PBMC3K | 10.87944414 | 0.008343052 | ** |
| CellTypeAI: Qwen3:32b vs CellTypist | PBMC3K | inf | 0 | *** |
| CellTypeAI: Qwen3:32b vs ScType | PBMC3K | inf | 0 | *** |
| CellTypeAI: Phi4:14b vs CellTypist | PBMC3K | 0 | 1 | ns |
| CellTypeAI: Phi4:14b vs ScType | PBMC3K | inf | 0 | *** |
| Gemini 2.5 Pro vs CellTypist | PBMC3K | 11.40266782 | 0.007603468 | ** |
| Gemini 2.5 Pro vs ScType | PBMC3K | 23.3826859 | 0.001823987 | ** |
| CellTypeAI: Qwen3:235b vs CellTypist | StrokePBMC | 1.010362971 | 0.418681641 | ns |
| CellTypeAI: Qwen3:235b vs ScType | StrokePBMC | 7.000372014 | 0.019801898 | * |
| CellTypeAI: Qwen3:32b vs CellTypist | StrokePBMC | -2.020725942 | 0.18071197 | ns |
| CellTypeAI: Qwen3:32b vs ScType | StrokePBMC | 9.959292144 | 0.009931966 | ** |
| CellTypeAI: Phi4:14b vs CellTypist | StrokePBMC | 1.732050808 | 0.225403331 | ns |
| CellTypeAI: Phi4:14b vs ScType | StrokePBMC | 8.577775428 | 0.013320035 | * |
| Gemini 2.5 Pro vs CellTypist | StrokePBMC | -0.757772228 | 0.527702159 | ns |
| Gemini 2.5 Pro vs ScType | StrokePBMC | 3.734734554 | 0.064802518 | ns |
| CellTypeAI: Qwen3:235b vs CellTypist | TS: Liver | inf | 0 | *** |
| CellTypeAI: Qwen3:235b vs ScType | TS: Liver | inf | 0 | *** |
| CellTypeAI: Qwen3:32b vs CellTypist | TS: Liver | inf | 0 | *** |
| CellTypeAI: Qwen3:32b vs ScType | TS: Liver | inf | 0 | *** |
| CellTypeAI: Phi4:14b vs CellTypist | TS: Liver | 6.423021745 | 0.023392201 | * |
| CellTypeAI: Phi4:14b vs ScType | TS: Liver | 24.24871131 | 0.001696354 | ** |
| Gemini 2.5 Pro vs CellTypist | TS: Liver | 18.35973856 | 0.002953518 | ** |
| Gemini 2.5 Pro vs ScType | TS: Liver | 46.88084186 | 0.000454687 | *** |
| CellTypeAI: Qwen3:235b vs CellTypist | TS: Lung | 25.78831202 | 0.001500293 | ** |
| CellTypeAI: Qwen3:235b vs ScType | TS: Lung | 74.8630849 | 0.000178381 | *** |
| CellTypeAI: Qwen3:32b vs CellTypist | TS: Lung | 13.38402897 | 0.005536153 | ** |
| CellTypeAI: Qwen3:32b vs ScType | TS: Lung | 53.53611587 | 0.000348722 | *** |
| CellTypeAI: Phi4:14b vs CellTypist | TS: Lung | 8.187876545 | 0.014590521 | * |
| CellTypeAI: Phi4:14b vs ScType | TS: Lung | 48.33996345 | 0.00042767 | *** |
| Gemini 2.5 Pro vs CellTypist | TS: Lung | 7.072540798 | 0.01941147 | * |
| Gemini 2.5 Pro vs ScType | TS: Lung | 25.47558063 | 0.001537268 | ** |
| CellTypeAI: Qwen3:235b vs CellTypist | TS: Small Intestine | inf | 0 | *** |
| CellTypeAI: Qwen3:235b vs ScType | TS: Small Intestine | inf | 0 | *** |
| CellTypeAI: Qwen3:32b vs CellTypist | TS: Small Intestine | 8.833459119 | 0.012574372 | * |
| CellTypeAI: Qwen3:32b vs ScType | TS: Small Intestine | 34.29460599 | 0.000849171 | *** |
| CellTypeAI: Phi4:14b vs CellTypist | TS: Small Intestine | 7.732369677 | 0.016317093 | * |
| CellTypeAI: Phi4:14b vs ScType | TS: Small Intestine | 25.91890316 | 0.001485246 | ** |
| Gemini 2.5 Pro vs CellTypist | TS: Small Intestine | 15.99599863 | 0.003885442 | ** |
| Gemini 2.5 Pro vs ScType | TS: Small Intestine | 45.95028907 | 0.000473277 | *** |
